# A host enzyme pseudouridylates an RNA bacteriophage genome and accelerates virus production

**DOI:** 10.64898/2026.10.15.759028

**Authors:** Janie S. Kim, Hossein Shenasa, Andrew Z. Fire, Nicole M. Martinez

## Abstract

RNA modifications are prevalent and potent regulators of gene expression across all domains of life. However, whether they exist in bacteriophages and whether they impact phage-host interactions are largely unexplored. To address this, we leveraged the ssRNA bacteriophage MS2 and its *E. coli* host. Using quantitative nanopore direct RNA sequencing and pseudouridine profiling, we found that essentially all copies of MS2 genomic RNA are pseudouridylated at a single position within the maturation protein gene. Through analysis of the surrounding sequence context, analysis of predicted structure, and genetic assignment of the pseudouridine to its installing enzyme, we demonstrate that the bacterial host enzyme RluA is responsible for pseudouridylation of the phage genome. In the absence of host RluA, the latent and eclipse periods are extended, indicating that the enzyme is important to early stages of phage infectivity. We find that this pseudouridine is present at varying occupancies in additional ssRNA coliphages, also installed by host RluA, and that the conserved features of RluA substrates are present in diverse ssRNA phages with host species beyond *E. coli*. Altogether, our findings identify RNA modification as a feature of bacteriophage genomes and establish host RNA-modifying enzymes as a class of infection-modulating phage host factors.

## Introduction

MS2 is a single-stranded, positive-sense RNA bacteriophage with a 3,569-nt genome that encodes only four proteins. The apparent simplicity belies the sophistication of its biology. Despite its status as an extensively studied model organism for over half a century, there remain numerous fundamental questions to be addressed about its biology and that of ssRNA phages more broadly. Although the genome of MS2 was the first to be fully sequenced^1–3^, a single base remained ambiguous across these early efforts. Furthermore, metatranscriptomics studies in recent years have additionally spurred a surge in newly identified RNA phages, whose diversity has ushered in an overhaul of their classification system and an increased interest in old and new unknowns of RNA phage biology^4–8^.

The minimalist genomes of ssRNA phages (typically <4.5 kb) impose unusual challenges in genetic regulation. With such limited coding capacity, translational regulation arises primarily from the genome’s own secondary and tertiary structure, and replication relies upon interactions between the phage genomic RNA and co-opted host translation factors^9–14,4,15,16^. The packaging of viral progeny involves similarly extensive RNA-protein contacts between an elaborately folded genome, a single copy of the phage’s maturation protein, and 89 coat protein dimers^17,18^. In other words, the replicative success of RNA phages like MS2 hinges upon the formation of *intra*molecular contacts within their own genomic RNA and upon *inter*molecular contacts with proteins originating from both host and phage. Chemical alterations to that genomic RNA may thus represent alterations to viral fitness.

Across all domains of life, RNA modifications have emerged as potent regulators of gene expression. These 170+ known noncanonical nucleosides alter the intrinsic biochemical and biophysical properties of an individual RNA molecule and, in turn, interactions of the RNA with itself and with other macromolecules^19–22^. Pseudouridine, one of the most abundant modifications, enhances base-stacking and stabilizes RNA duplexes^23–25^, decreases RNA backbone flexibility^23,26^, protects against cleavage by both auto-hydrolysis^27^ and nuclease activity^28,29^, and alters the binding of proteins to RNA^26,30–33^. Such changes impact gene expression at the co-and post-transcriptional levels by altering the interactions of an RNA molecule within both endogenous and exogenous environs^24,34^, and they are the basis for suppressing the activation of eukaryotic innate immune sensor responses by modified compared to unmodified RNAs of endogenous, bacterial, and synthetic origins^19,35,30^.

Despite this gamut of molecular impacts, it is unknown whether RNA modifications intersect with the life cycles of RNA phages^36^. Much of the attention directed towards RNA modifications in recent years has been centered on eukaryotes, with a smaller fraction towards bacteria and a vanishingly smaller subset towards their viruses. With recent surges in discovery of diverse RNA phages^4,6,7^, of molecular and biological consequences of RNA modifications^19–22^, and of molecular subterfuge in the phage-host “arms race” involving ribonucleases^37–41^, this convergence of RNA biology and microbiology presents a timely question: What is the relevance of modified ribonucleosides in the biology of RNA phages?

In recent years, various RNA modifications in RNAs beyond tRNAs and rRNAs and their installing enzymes have been catalogued across multiple bacterial species^42–46^. These modifications and their installing enzymes impact bacterial physiology and fitness, as has been demonstrated with a few isolated modifications^47,48^ and with several enzymes^49–55^. This expanding bacterial corpus leaves a phage-shaped question in its wake: Do RNA phages, which replicate within that same bacterial cytoplasmic milieu, also harbor RNA modifications, and if so, do these modifications have fitness-relevant roles? With the precedent set by mammalian viruses, in which RNA modifications installed by host enzymes alter viral replication and infectivity^56,57^, we first turned our attention to a particularly tractable phage-host model, MS2 phage and *E. coli*.

## Results

### The genome of the RNA bacteriophage MS2 is modified

To determine whether the MS2 genomic RNA harbors modified nucleotides, we first performed long-read Oxford Nanopore Technologies (ONT) direct RNA sequencing as an initial survey of the whole 3,569-nt genome. In standard ONT direct RNA sequencing workflows, threading RNA through a nanopore with an applied voltage allows diverse modifications to be observed by deviations in ionic current, dwell time, and base miscalls from those of the four canonical bases (Fig. 1A and 1B). Sequencing MS2 genomic RNA and extracting all positions with ≥20% U-to-C base miscalls^58^ nominated several putative modification sites (Fig. 1C, Fig. S1A-C, Tables S3 and S4). These sites had high proportions of U-to-C base miscalls indicative of pseudouridine (Fig. 1B and 1C, Table S3 and S4). This single-nucleotide-resolution signature of pseudouridine has been benchmarked extensively with genetic assignment to pseudouridine synthases and with modified and unmodified synthetic controls encompassing varying sequence contexts^59–61,58^. It was possible that U-to-C miscalls might represent true sequence diversity in the pool of RNAs at these positions; further sequencing of the MS2 RNA with modification-insensitive cDNA Illumina sequencing ruled out this possibility (Fig. 1C). Basecalling with ONT’s pseudouridine-aware Dorado basecaller model further supported position 924 as a putative pseudouridine (Table S5).

**Figure 1.**
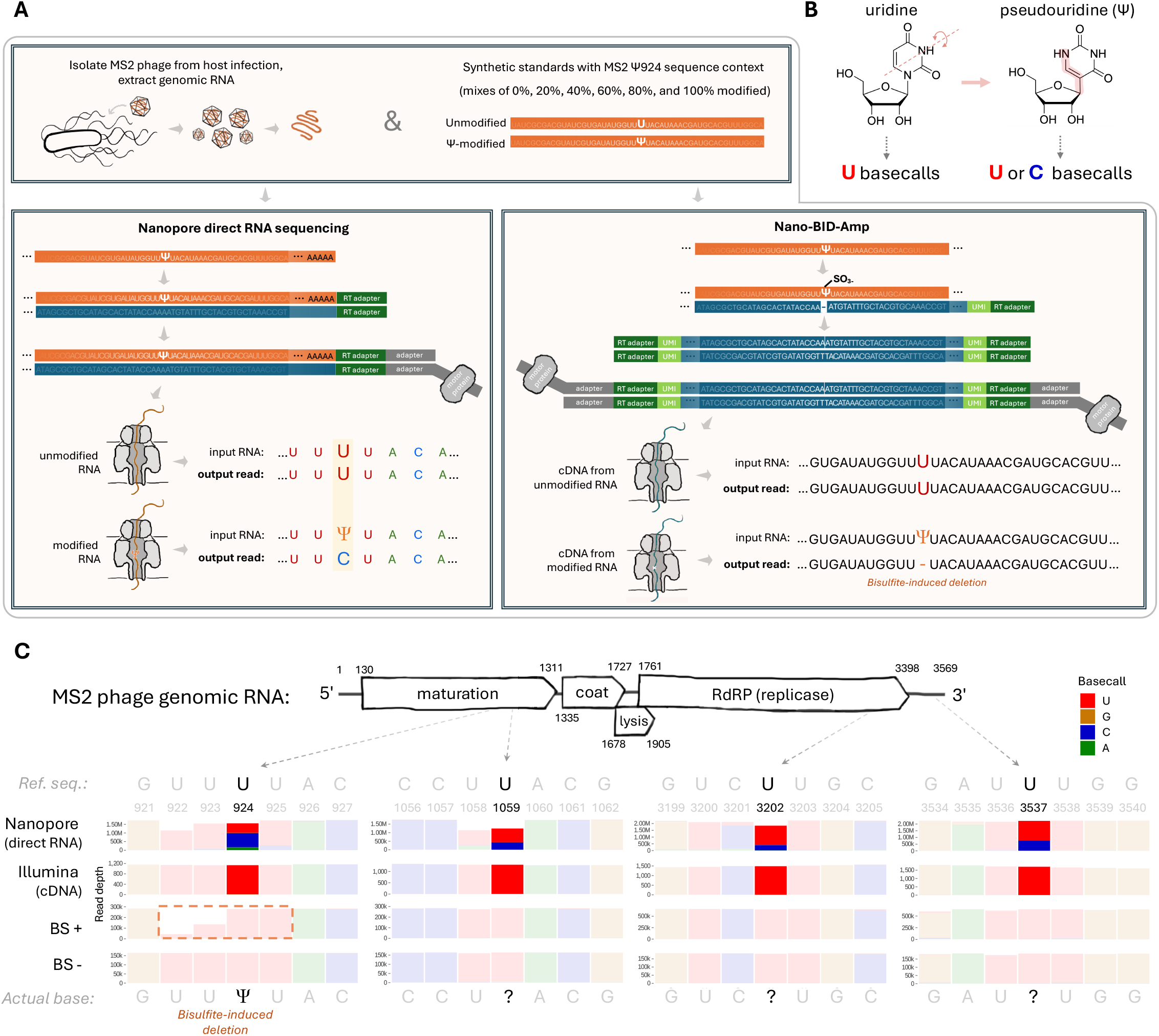
The genome of the RNA bacteriophage MS2 is modified. (A) Sequencing workflow for identification of modifications in MS2 phage genomic RNA. Mixtures of modified and unmodified synthetic standards at different ratios were included in the same workflow to generate a calibration curve for determining stoichiometry. Oxford Nanopore Technologies (ONT) direct RNA sequencing library prep involves reverse-transcription of the RNA and ligation of the motor protein adapter, followed by sequencing of the RNA strand, from which modifications like pseudouridine can be identified by base miscall signatures. Nano-BID-Amp library prep involves treating the RNA with bisulfite, which results in deletions at pseudouridine sites when the RNA is reverse-transcribed and sequenced. **(B)** The chemical structure of pseudouridine, an isomer of uridine. Pseudouridine sites exhibit characteristic U-to-C base miscalls in ONT direct RNA sequencing reads. **(C)** Positions across the MS2 phage genomic RNA with U-to-C base miscalls, as identified with ONT direct RNA sequencing (n=3), that are not genomic polymorphisms, as verified with Illumina cDNA sequencing (n=2). All sites were subjected to bisulfite-induced deletion and sequencing as orthogonal validation for candidate pseudouridine sites (n=1; “BS+” indicates samples treated with bisulfite and “BS-“ indicates untreated control samples). Dashed orange box indicates ambiguity in deletion assignment between the four consecutive uridines in positions 922-925 (read alignment by default assigns deletions to the 5’-most position in U-tracts). Y-axis values indicate read depth. Genomic coordinates shown are from RefSeq and include stop codons.

To orthogonally assess potential pseudouridine sites, we performed bisulfite-induced deletion (BID) and sequencing. Bisulfite-based detection hinges upon the formation of a ring-opened adduct when a pseudouridine selectively reacts with bisulfite at neutral pH; such adducts manifest as characteristic deletions when the RNA is reverse-transcribed into cDNA and sequenced^62–64,58^ (Fig. 1A). We therefore applied our targeted Nano-BID-Amp workflow that relies on this principle^42^ to the MS2 genomic RNA (Fig. 1A). This yielded the pseudouridine-specific deletion signature only around position 924 (Fig. 1C, Fig. S1D).

Although bisulfite-induced deletion typically manifests as the deletion of one base, at our MS2 site we observed both single and double deletions (Fig. 1C). This can be attributed to reverse transcriptase behavior: polymerase slippage on homopolymeric tracts has long been known to produce base deletions in a manner dependent on surrounding sequence context^65,66^, and mutational adducts including ring-opened sites have been previously observed to produce penetrant double-deletion signatures^67,68,62^. Both single-deletions and double-deletions are arbitrarily aligned to the 5’-most U out of the four-U tract, as alignment is unable to unambiguously assign a deletion to any particular U within a multiple-U motif^64,69^.

Together, nanopore direct RNA sequencing and Nano-BID-Amp confirm the presence of a modification consistent with pseudouridine at genomic position 924 within the phage’s maturation protein gene.

### Host enzyme RluA pseudouridylates MS2 RNA genome at high stoichiometry

The compact 3.5 kb MS2 genome encodes only four proteins—the maturation protein, which mediates host attachment, genome entry, and virion assembly^70^; the coat protein, which comprises the bulk of the capsid^18^; the lysis protein, which mediates host lysis by disrupting the cell membrane^71,72^; and the RNA-dependent RNA polymerase (RdRp), which synthesizes genomic copies^3,16^ (Fig. 2A). We found no unannotated pseudouridine-synthase-like domains within these genes, indicating that Ψ924 likely does not arise from an enzyme endogenous to the phage (see Methods). We therefore turned our attention to host enzymes.

**Figure 2.**
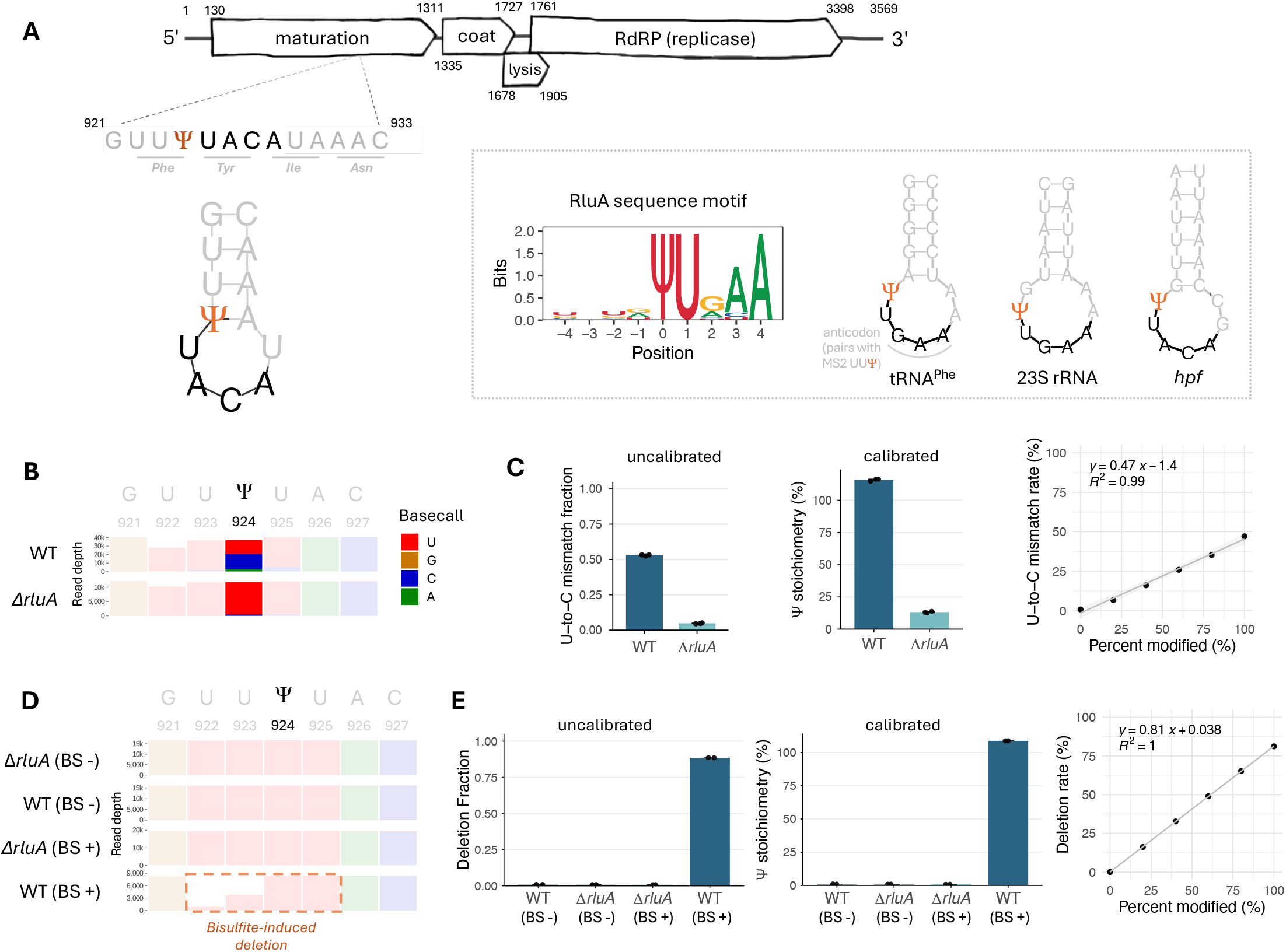
Host enzyme RluA pseudouridylates MS2 RNA genome at high stoichiometry.(A) The sequence, predicted secondary structure, and coordinates of the RluA-modified stem-loop in MS2 genomic RNA. RNAfold with a folding window of ±50 nt was used to model the structure (see Methods). Inset, from left to right: RluA recognition sequence logo, generated from the known four tRNA substrates, 23S rRNA substrate, and 31 high-confidence mRNA substrates from published studies; tRNA, rRNA, and mRNA examples of known RluA substrates from the *E. coli* transcriptome^75,76,43^. **(B)** Representative ONT direct RNA sequencing basecall profile of MS2 genomic position 924 and flanking ±3 bases for MS2 from both wild-type host and Δ*rluA* host (n = 3). Y-axis values indicate read depth. **(C)** U-to-C mismatch fractions from ONT direct RNA sequencing of independent biological replicates, for MS2 isolated from wild-type and Δ*rluA* host infections (n=3). fi stoichiometry at position 924 in both wild-type and Δ*rluA* MS2 was calculated by fitting a linear line of best fit to the synthetic standard calibration curve. The 0%-modified and 100%-modified synthetic standard reads were computationally mixed to generate a calibration curve. Gray shading represents 95% confidence intervals. **(D)** Representative Nano-BID-Amp trace of MS2 genomic position 924 and flanking ±3 bases, for MS2 isolated from wild-type and Δ*rluA* host infections (n = 2; “BS+” indicates samples treated with bisulfite and “BS-“ indicates untreated control samples). Dashed orange box indicates ambiguity in deletion assignment between the four consecutive uridines in positions 922-925 (read alignment by default assigns deletions to the 5’-most position in U-tracts). Y-axis values indicate read depth. **(E)** Deletion fractions from Nano-BID-Amp for MS2 from both wild-type host and Δ*rluA* host (n=2; “BS+” indicates samples treated with bisulfite and “BS-“ indicates untreated control samples). fi stoichiometry at position 924 in both wild-type and Δ*rluA* MS2 was calculated by fitting a linear line of best fit to the synthetic standard calibration curve. The 0%-modified and 100%-modified synthetic standard reads were computationally mixed to generate a calibration curve. Gray shading represents 95% confidence intervals.

Pseudouridine synthases recognize their targets through RNA sequence and/or structural features^73,74,43,58^. Analysis of the sequence and predicted secondary structure surrounding the Ψ924 site revealed a resemblance to the known recognition motif of the *E. coli* pseudouridine synthase RluA (Fig. 2A). The motif for canonical substrates of RluA—fi32 in the anticodon stem-loops of four tRNAs and fi746 on the 23S rRNA^75,76^—comprises a UURAA located within a short stem-loop, in which the underlined U is pseudouridylated and is located in the loop near the 5’ side of the stem^43^ (Fig. 2A and Table S6; “R” indicates either A or G). The genomic context in which MS2 Ψ924 is situated is a close but not exact match: UUACA, with the pseudouridylated first U in the closing base-pair of the stem and UACA in the loop region (Fig. 2A). Given that the closing base-pair of stem-loops is often prone to fraying, especially U·A pairs^77^, we suspected that this MS2 site could be a bona fide target of RluA. We were additionally encouraged by a recent expansion of the known modification repertoire of *E. coli* RluA to various mRNA substrates^43^; combining all canonical targets and these 31 newly identified mRNA targets yielded a recognition motif logo that gave further credence to our hypothesis (Fig. 2A and Table S6).

We therefore evaluated the modification status of MS2 in the absence of host RluA. We infected Δ*rluA* and wild-type host strains with MS2 in double-layer agar plaque assays. When we extracted RNA from the resulting phage progeny and performed ONT direct RNA sequencing and Nano-BID-Amp, we observed a loss of Ψ924 in phage from Δ*rluA* host infection but not in the wild-type counterpart (Fig. 2B-E; Fig. S2A and S2B; Table S3). This indicated that *E. coli* RluA is responsible for the modification at position 924 and solidified the model that this modification is a pseudouridine. When we extracted all genomic sites with a ≥20% difference in base miscalls between wild-type reads and Δ*rluA* reads in three replicates, the only pseudouridine was position 924 (Table S7).

Because the stoichiometry of a modification will constrain any biological function, we determined the stoichiometry at position 924 using synthetic standards. These standards were RNA oligos encompassing 25 bases upstream and 24 bases downstream of Ψ924 in the MS2 genome, with or without a pseudouridine at the site corresponding to genomic position 924 (Table S2). By mixing these modified or unmodified standards at various ratios from 0% modified to 100% modified and including them in both our ONT direct RNA sequencing and Nano-BID-Amp workflows (Fig. 1A), we were able to obtain both physically mixed and computationally mixed calibration curves to determine modification stoichiometry from raw signals (Fig. 2C and 2E; Fig. S2C-G; Table S3 and S5). These standards with a single pseudouridines exhibit the same mismatch profile (e.g., U-to-C mismatches in ONT direct RNA sequencing and double-deletions in Nano-BID-Amp) as the MS2 viral genome (Fig. S2E; Table S3 and S5). We indeed observed a concordance between the stoichiometry estimates obtained from both approaches, in which these values were slightly overestimated but the difference between wild-type and Δ*rluA* was consistently ∼100 percentage points (Fig. 2C and 2E). Thus, both orthogonal approaches converge upon stoichiometric or near-stoichiometric pseudouridylation at position 924 in a manner highly dependent upon RluA. It is also notable that a 100%-modified synthetic standard produced direct RNA U-to-C miscalls only ∼45% of the time and bisulfite-induced deletions 84% of the time, which underscores the impact of context-dependent effects and the value of synthetic-standard-based calibration in estimating stoichiometries^78,79,63,61,58^.

### Loss of host RluA extends the viral latent period

Given that virtually every MS2 genome copy harbors a pseudouridine, we wondered whether loss of this modification could affect viral fitness. We therefore sought to compare the infectivity of MS2 phage on wild-type and *ΔrluA* hosts. We observed in double-layer agar plaque assays^80^ that MS2 forms plaques on both hosts (Fig. 3A), indicating that host RluA is not essential to phage infectivity. When we compared the efficiency of plaquing (EOP) on the Δ*rluA* host relative to that on the wild-type host, we did not observe a significant difference, although a slight trend of lesser EOP on the mutant may be present (Fig. 3A and 3B). We note that plaque assays are an endpoint readout for infection and do not capture the finer-grained dynamics of phage infection leading up to that point.

**Figure 3.**
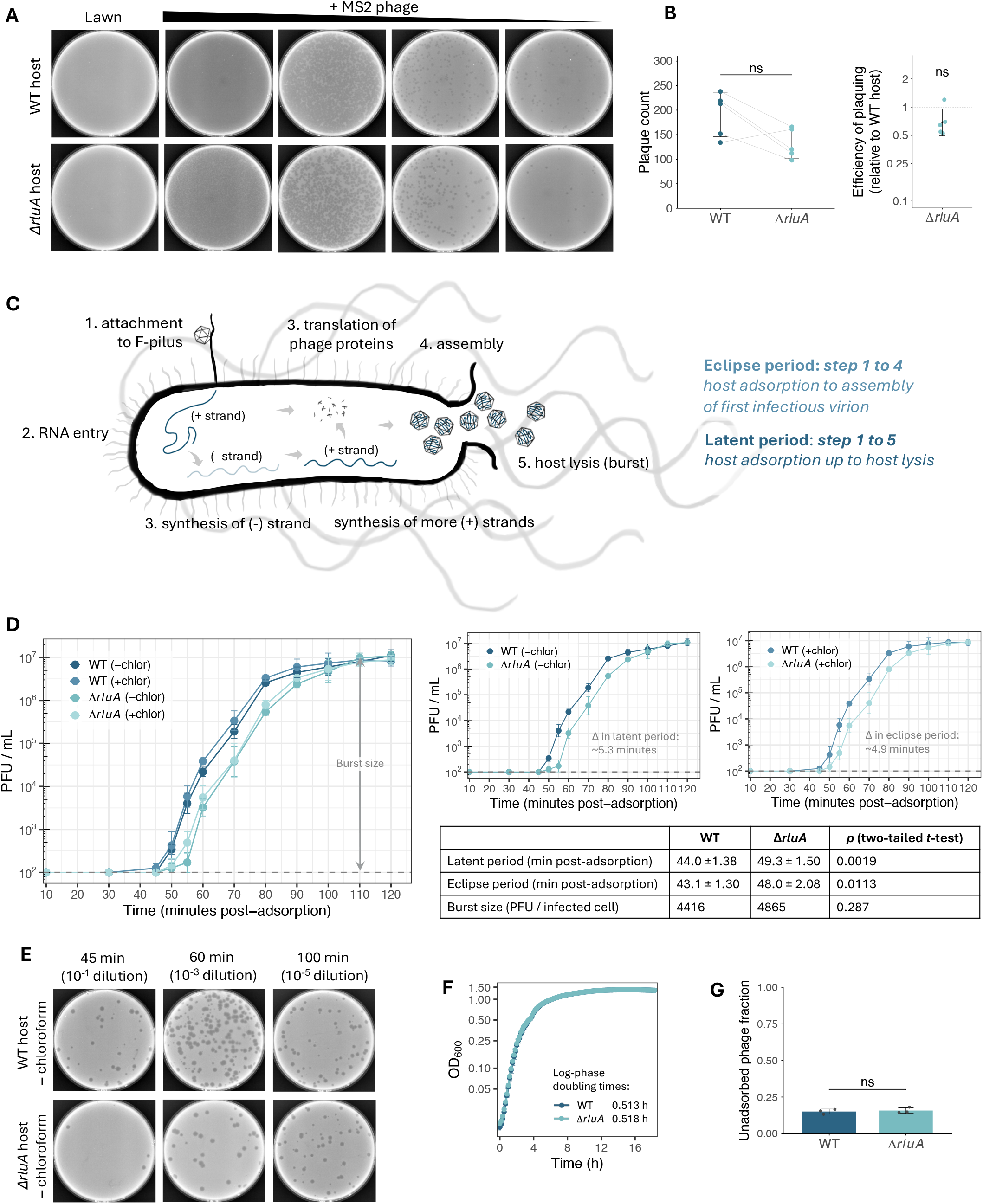
Loss of host RluA extends the viral latent period. **(A)** Representative images of double-layer agar plaque assay comparing MS2 phage efficiency of plaquing on wild-type host versus Δ*rluA* host (n=5). **(B)** Left: Raw plaque counts on wild-type and Δ*rluA* hosts. Bars represent the mean ± SD, individual points represent biological replicates, gray lines indicate wild-type-and-Δ*rluA* matched pairs (n = 5 pairs; two-tailed paired *t-*test, *p* = 0.066). Right: Efficiency of plaquing (EOP) on Δ*rluA* host relative to wild-type host. Each raw Δ*rluA* plaque count was divided by its matching raw wild-type plaque count. Dashed line is the wild-type reference (EOP = 1). Individual points represent biological replicates. The error bar range indicates ×/÷ geometric SD and the central black point indicates the mean. (n = 5 pairs; two-tailed paired *t-*test on log_10_-transformed values, *p* = 0.066). **(C)** Schematic of MS2 infectious life cycle, steps numbered in chronological order. The latent period spans from adsorption (step 1) up to host lysis (step 5). The eclipse period is a subset of the latent period, spanning from adsorption to the formation of the first infectious virion in the cytoplasm. **(D)** One-step phage growth assay for wild-type MS2 on wild-type and Δ*rluA* hosts at MOI = 0.01. Datapoints represent the geometric mean of three biological replicates, with error bars indicating ×/÷ geometric SEM. Dashed gray line indicates assay limit of detection (1 plaque). Leftmost plot depicts all strains ± chloroform; smaller plots on the right depict-chloroform and +chloroform data only. Estimates of eclipse period, minimum latent period, and burst size are provided in the table (see Methods). **(E)** Representative plaque assay plates from key one-step growth curve timepoints—45, 60, and 100 minutes—plated at the same indicated dilutions for both wild-type and *ΔrluA* host samples without chloroform-induced lysis. **(F)** Growth curves of wild-type and Δ*rluA* hosts. The datapoints represent the average of three biological replicates ± SEM. Log-phase doubling times were estimated from a linear line of best fit generated from datapoints between 0.5 and 2.5 hours. The difference in doubling times is not statistically significant (Welch’s *t-*test, *p-*value = 0.89). **(G)** Fraction of unadsorbed phage relative to input PFU, following adsorption at MOI = 0.01 for 10 minutes at 37 °C. Bars represent the mean ± SD. Welch’s *t*-test, *p-*value = 0.68.

To obtain insight into the earlier stages of the phage life cycle (Fig. 3C), we performed standard one-step phage growth curves^81,82^. We infected log-phase host cells with MS2 phage, washed away unadsorbed phage, and then diluted these cultures 1,000-fold to preclude secondary infections. Enumerating the plaque-forming unit (PFU) counts of these cultures over the course of two hours allowed us to compare a single synchronized infectious cycle of MS2 between the two host strains. By enumerating PFU of all timepoints with or without chloroform-induced artificial host lysis (Fig. 3D and 3E; Fig. S3), we obtained estimates for both extracellular phage and total phage, which allowed us to compare the following between wild-type and Δ*rluA* hosts: 1) the latent period, the time spanning host adsorption up to host lysis; 2) the eclipse period, the subset of the latent period defined as the time during which phage RNA and proteins are being synthesized inside the host cell (spanning host adsorption to intracellular assembly of an infectious phage particle); and 3) the burst size, the number of infective phage progeny produced per host cell (Fig. 3C).

By comparing the timepoints without artificially induced host lysis, we estimated that the latent period is ∼5 minutes longer in MS2 infecting Δ*rluA* host (∼49 minutes) than wild-type host (∼44 minutes) (Fig. 3D and 3E). This delay manifests as a consistent ∼10-fold difference in PFU between wild-type and Δ*rluA* hosts throughout the rise phase, until both phage populations plateau at a final PFU per infected host cell of 4 x 10^3^ to 5 x 10^3^, with no significant difference in burst size (Fig. 3D and 3E). This effect could not be attributed to a simple growth defect in the Δ*rluA* host, as the doubling times of both strains in liquid culture were not significantly different (mean exponential doubling time of 0.513 hours and 0.518 hours for wild-type and Δ*rluA* hosts, respectively; Fig. 3F). Given that the latent period is longer for Δ*rluA* but the burst size does not display a compensatory increase, we suspected that the mechanism underlying the defect occurs in a step of phage infection preceding host cell lysis (i.e., there is not a build-up of phage progeny released once the host cell is finally lysed after a delay).

Indeed, by comparing the timepoints with artificially induced host lysis, we estimated that the eclipse period is also ∼5 minutes longer in Δ*rluA* host infection than wild-type host infection (Fig. 3D). This extended eclipse period supports the longer latent period being not primarily attributable to a defect in host cell lysis but rather to an earlier step of infection. When we compared MS2 adsorption efficiency on wild-type and Δ*rluA* hosts, we found no statistically significant difference (Fig. 3G). This indicated that loss of host RluA does not produce defects in the F-pilus or other cell surface characteristics in a way that affects wild-type MS2 adsorption.

Altogether, we deduce that the latent period defect when MS2 infects a Δ*rluA* host arises from an intracellular process following adsorption but preceding host cell lysis.

### Pseudouridylation by host RluA is conserved in close MS2 relatives

Several RNA modifications are evolutionarily conserved across *Bacteria*^21,46^. Given the presence of the MS2 Ψ924, we wondered how broad this phage pseudouridylation could be beyond the case of MS2. The local structural context in which a primary sequence motif is embedded has been demonstrated to be a key determinant of modification efficiency for several pseudouridine synthases^44,45,17,31^. In the case of RluA, substrates fold into stem-loops, as supported by DMS-informed structure prediction^43^. In order to perform a wider search for RluA-modified sites across other ssRNA phages, we first inspected all known terminal stem-loop substrates of RluA with experimentally determined structures to define loop and stem lengths compatible with modification (Fig. S4A) ^75,76,43^ (n = 21; Table S8). We also noticed a consistent trend in the position of the UURHA primary motif within the loop of substrates, where the motif is situated such that there are always fewer loop bases 5’ of it than 3’ of it, irrespective of loop length (Fig. S4A; Table S8; “R” indicates A/G and “H” indicates A/C/U). We used these observations to define and computationally search for RluA substrate stem-loops (see Methods).

We first applied these criteria to computationally folded (RNALfold) genome sequences of other classic “Group I” ssRNA coliphages, which range between approximately 77.5% to 98.9% sequence identity to MS2 (Fig. 4A; Table S9 and S10). (“Group I” coliphages were designated strains of *Emesvirus zinderi* under the 2021 ICTV reclassification^8^.) All of these phages—R17, fr, M12, JP501, DL1, DL16, J20, ST4, DL52, DL54—encode the RluA substrate stem-loop within the same homologous region of their maturation protein genes (Fig. 4A, Table S11). The only phage lacking the exact MS2 sequence was JP501: UUAUA instead of UUACA, a single nucleotide difference that still meets RluA-modification criteria (Fig. 2A).

**Figure 4.**
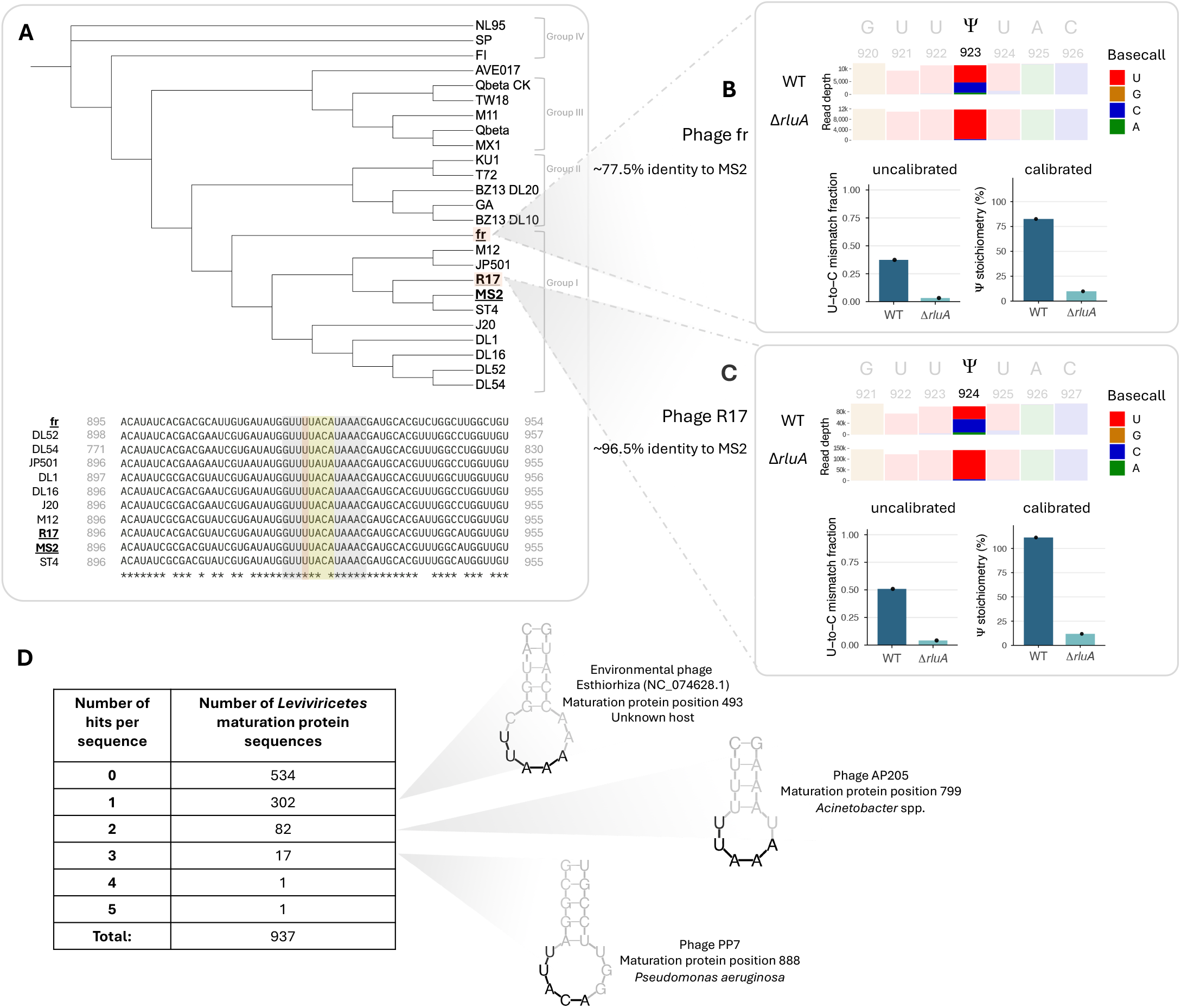
Pseudouridylation by host RluA is conserved in close MS2 relatives. **(A)** Top: Minimum evolution phylogenetic tree for MS2 and other ssRNA coliphages (pairwise distance between genome sequences). All sequences—with the exception of “Qbeta CK,” which was a gift from the lab of Charlotte Knudsen—were obtained from RefSeq and GenBank (Table S9). Bottom: Nucleotide alignment of strains of *Emesvirus zinderi* (“Group I” coliphages): 60-nt window containing the RluA stem-loop (gray), RluA recognition sequence motif (yellow), and position pseudouridylated in MS2 (orange). **(B)** Top: ONT direct RNA sequencing basecall profile for phage fr from wild-type and *ΔrluA* hosts (n=1), depicting the pseudouridylated site and flanking bases. Y-axis values indicate read depth. Bottom: Raw uncalibrated U-to-C mismatch fractions at position 924 and calibrated pseudouridylation stoichiometries derived using the calibration curve from the MS2 synthetic standards (Fig. 2C; Fig. S4F). **(C)** Top: ONT direct RNA sequencing basecall profile for phage R17 from wild-type and *ΔrluA* hosts (n=1), depicting the pseudouridylated site and flanking bases. Y-axis values indicate read depth. Bottom: Raw uncalibrated U-to-C mismatch fractions at position 924 and calibrated pseudouridylation stoichiometries derived using the calibration curve from the MS2 synthetic standards (Fig. 2C; Fig. S4F). **(D)** *In silico* search for putative RluA-modified stem-loops across 937 *Leviviricetes* maturation protein sequences yielded 403 sequences with hits. The distribution of hits per sequence is listed in the table. Putative RluA stem-loops from three example phages with 1 hit, 2 hits, or 3 hits in their maturation protein sequences are shown, with the RluA primary sequence motif in black. “Position” indicates position of the first U in the UURHA motif. Detailed information on all *in silico* hits can be found in Table S12.

Conversely, this motif was absent in the homologous maturation protein region (see Methods) of more distantly related *Emesvirus* coliphages: *Emesvirus japonicum* strains GA, KU1, T72, and BZ13 isolates DL10 and DL20 (historically classified as “Group II” coliphages; ∼56-59% identity to MS2) and AVE017, the type strain of *Emesvirus piscicola* (∼55% identity to MS2; Tables S9-11). The motif was also absent in the homologous maturation protein regions of more distantly related *Qubevirus* coliphages: *Qubevirus durum* strains Qβ, M11, MX1, TW18 (historically classified as “Group III” coliphages; ∼43-44% identity to MS2) and *Qubevirus faecium* strains FI, SP, and NL95 (historically classified as “Group IV” coliphages; ∼44-45% identity to MS2; Tables S9-11). This suggested that the RluA-installed pseudouridine may be a feature shared by *Emesvirus zinderi* strains.

To test the hypothesis that other ssRNA coliphages closely related to MS2 are modified by host RluA, we performed ONT direct RNA sequencing on a selection of additional coliphages. In the closely related phages R17 (∼96.5% identity to MS2) and fr (∼77.5% identity to MS2), we indeed observed pseudouridylation at the same homologous position as in MS2. These phages also have high estimated modification stoichiometries of ∼83% for fr and of ∼100% for R17 (Fig. 4B and 4C; Fig. S4D-G; Tables S3, S5, S7). For both phages, the pseudouridine is absent in virions isolated from Δ*rluA* host infection (Fig. 4B and 4C), indicating that the host RluA is the responsible pseudouridine synthase. Although our *in silico* pipeline for identifying putative RluA substrates yielded three additional sites in phage fr conforming to the RluA motif as defined, we found no evidence of modification at these sites (Table S11, S7). In the more distantly related phage Qβ (a strain from the lab of Charlotte Knudsen; ∼43% identity to MS2), we found no evidence of pseudouridine (Fig. S4B, C, G; Table S7).

To survey for the possibility of modification in phages beyond the classic lab-cultivable coliphages, we expanded our computational search to the taxonomic class *Leviviricetes,* which comprises all non-enveloped, icosahedral, positive-strand ssRNA phages that infect different host bacterial species via adsorption to various pili^5^. We computationally folded 937 *Leviviricetes* maturation protein coding sequences from NCBI Virus and searched these predictions for putative RluA stem-loops. 534 sequences had no predicted stem-loops, while the remaining 403 averaged 1.67 stem-loops per sequence (Fig. 4D; Fig. S4A and S4H; Table S12). Although the majority of these phages were identified in metatranscriptomics studies and therefore remain uncultivated with unknown hosts, a few stem-loops were predicted in phages with known host species beyond *E. coli* that genomically encode RluA (Fig. 4D and Table S12).

## Discussion

### An RNA phage is modified at high stoichiometry by a host pseudouridine synthase

The RNA modification landscape across mRNAs has been sparsely characterized to date in microbes^42,43^. Here, we discovered a modification in a bacteriophage’s genomic RNA installed by a bacterial host enzyme. This pseudouridine at genomic position 924 was independently reported by two concurrent studies while this work was in progress using ONT direct RNA sequencing^83,84^ and the presence of a pseudouridine in MS2 was corroborated by bulk nucleoside mass spectrometry^84^. We further ascertain that this Ψ924 is present in virtually every phage RNA molecule and is installed by the *E. coli* pseudouridine synthase RluA, expanding the enzyme’s range of substrates beyond endogenous rRNA, tRNAs, and mRNAs to exogenous phage RNA. Notably, position 924 corresponds to the single base that was missing in initial sequencing efforts in the early 1970s^1^ and later inferred to be a uridine through a process that is unclear^2,3^. Thus, this mystery persisted for fifty years.

This high-stoichiometry MS2 phage Ψ924 is unusual. Although we could not find reports of RNA modifications in other RNA bacteriophages, we found a limited sample of reported stoichiometries in eukaryotic RNA viruses, none of which were estimated full-occupancy: ∼20% for an m^5^C site in dengue virus and three m^6^A sites in the 3’ UTR region of HIV-1 with estimated occupancies of 47%, 79%, and 89%^85^. Stoichiometric data is sparse given that the majority of reports of modification use non-quantitative profiling methods^86^. The identity of our orthogonally validated MS2 modification as a pseudouridine was also notable, as pseudouridines have not been unequivocally detected to date in other RNA viruses and reports of modification have conflicted^87–90^. The stoichiometry of a modification is important because it constrains the mechanisms through which the modification can exert biological impacts. A modification residing on every viral transcript acts broadly: if Ψ924 confers resistance to host RNase activity, this applies to the bulk population; if it architects a particular higher-order structure, this is true for every genome copy; if it alters translation efficiency of the gene it is situated within, this is relevant each time that gene is translated. By contrast, sub-stoichiometric modifications could represent fine-tunable regulation or become amplified as in the case of selective packaging^91^.

### Ψ924 resides in a genomic position that could plausibly impact translation and packaging

Ψ924 is located in the wobble position of the Phe265 codon in the maturation protein (Fig. 2A), and pseudouridines can impact translation efficiency^92,35,34,93^. Translation of this protein—which primarily proceeds in the nascent RNA before the formation of higher order structures in the mature RNA occlude ribosome access^2,12^—may also be impacted if the pseudouridine influences the folding kinetics. There is also potential for translational interplay between modified phage RNA and modified host RNA: *E. coli* tRNA^Phe^ is itself a substrate of RluA and is pseudouridylated two bases 5’ of the anticodon in a site known to impact translational fidelity and codon-anticodon interaction^3,63^ (Fig. 2A), raising the possibility of translational cooperation between the phage’s pseudouridylated Phe and the host’s pseudouridylated tRNA^Phe^. There is also a precedent for RNA modification impacting viral packaging^91^. In MS2, Ψ924 lies between two cryo-EM-verified packaging signal stem-loops^95,96^. The sequences of inter-packaging-signal regions have been implicated in packaging selectivity^97^, raising the possibility that the pseudouridine here could alter local rigidity and influence folding and packaging.

Pseudouridine may also alter susceptibility to host RNases, in an echo of DNA phages classically installing DNA modifications to prevent cleavage by host restriction enzymes. It is possible that an analogous principle of bacterial innate immune evasion applies to RNA phages and RNA modifications, given that known phage defense systems cleave various RNA targets^98–100,37,39,40^ and that pseudouridines inhibit cleavage by several RNases^30,28,29^. There is indeed a correlation between pseudouridylation status and transcript abundance in various bacteria^44,46^. Given the comparatively small genomes of RNA viruses compared to DNA viruses^101^, which encode their own DNA modification enzymes, “co-opting” the host enzyme could represent an RNA virus strategy to bypass intrinsic gene-encoding limitations and ensure their own replicative success.

### A bacterial pseudouridine synthase promotes phage intracellular replication post-adsorption but pre-lysis

Host RluA augments the early-stage infectivity of MS2 phage. The conceivable fitness consequence in the absence of RluA—a 5-minute delay in host cell lysis (Fig. 3D)—is nontrivial. In the context of phage population dynamics, where burst sizes are large and generation times are short, five minutes may be a deciding factor in which phage variant establishes itself by outcompeting others: By the 220-minute mark, wild-type-host MS2 with the shorter latent period will have completed ∼5 rounds of infection, while Δ*rluA*-host MS2 with the longer latent period will only have completed ∼4 rounds of infection, which already approximates to a fold difference in PFU of one extra burst size, roughly 1,000-to 10,000-fold. Another study reported that DNA phage T5 infection of *E. coli* lacking m^6^A enzymes compared to wild-type host exhibits a similar ∼5-minute delay in the rise period but also a decrease in burst size^53^. Conversely, in human hepatitis C virus infection, m^6^A restrains viral infectivity; depleting the host’s methyltransferase increases infectious particle production^56^. These results indicate that RNA modifications can have context-dependent phenotypes that are unique to the enzyme, host, and virus.

Because *E. coli* RluA installs pseudouridine onto a broad range of endogenous RNA substrates spanning tRNA to rRNA to mRNA, it is possible that loss of these native host modifications could pleiotropically affect infectivity by altering expression of host RNA or proteins relevant to phage infection. We ruled out an impact on exponential doubling time (Fig. 3F) or cell surface changes that impact adsorption (Fig. 3G). One remaining possibility is that loss of RluA-installed pseudouridines on endogenous RNAs alters efficacy of abortive infection; a plethora of phage defense systems trigger abortive infection via nucleases that target their own tRNAs, rRNAs, and mRNAs^38,41,102,103^, whose stability and structure are known to be affected by modifications^94^.Regardless of precise mechanism, these findings uncover a role for pseudouridine synthases in phage-host dynamics.

### Pseudouridylation by host RluA is conserved in MS2 and related phages

We found that the genomes of multiple ssRNA coliphages are pseudouridylated and that the features shared by substrates of host enzyme RluA are present among diverse ssRNA phages with hosts beyond *E. coli*, suggesting that RNA modification may be a broader phenomenon. That the RluA primary motif is identical between phages MS2, R17, and fr yet pseudouridylation stoichiometries differ between their genomes (Fig. 4A) suggests that broader sequence context (Fig. S4F) or differing host cellular conditions during infection by different phages may impact host RluA activity. As metatranscriptomic and other studies rapidly expand the known diversity of RNA phages, determining the prevalence and functional impacts of modification in phage genomes will be valuable for understanding epitranscriptomic mechanisms in phage-host interactions.

Seventy-five years after the discovery of pseudouridine as the first modified nucleoside (1951) and fifty years after the publication of the MS2 genome as the first complete genome sequence (1976), we identified within this genome a host-installed pseudouridine. Altogether, we characterized the stoichiometry, biological impact of host RluA, and conservation of the RluA-installed pseudouridine in the genomic RNA of MS2 and related ssRNA phages, thereby revealing a new layer of complexity within ssRNA phages and a hitherto unrecognized role of RNA modification enzymes in phage-host dynamics.

## Supplementary Figures

**Supplementary Figure 1.**
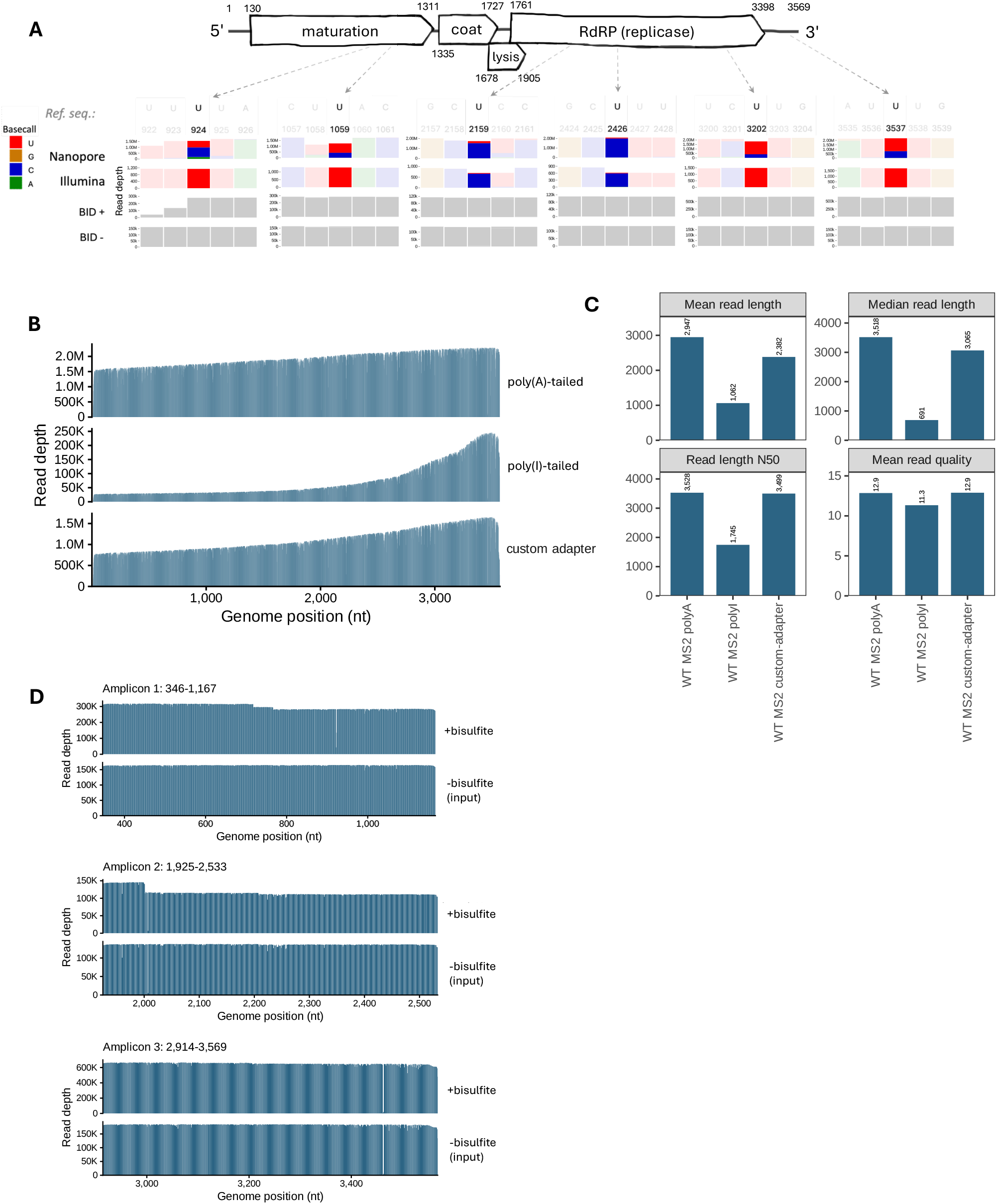
Sequencing coverage across MS2 genomic RNA. **(A)** All sites in MS2 RNA obtained from Roche with U-to-C base miscalls against the reference sequence, as identified with ONT direct RNA sequencing (n=3), including two extra sites identified as SNPs with Illumina cDNA sequencing (n=2). All sites were subjected to bisulfite-induced deletion and sequencing (n=1) as orthogonal assessment of putative pseudouridine sites. Y-axis values indicate read depth. Genomic coordinates shown are from RefSeq and include stop codons. **(B)** Per-base coverage of ONT direct RNA sequencing of MS2 RNA obtained from Roche. Three different library prep methods were used for each of the three runs (poly(A)-tailing, poly(I)-tailing, and custom adapter with complementarity to the last 10 3’-most bases of the MS2 genome). Representative data shown in Fig. 1C and S1A are with the poly(A)-tailed sample. **(C)** Run statistics for ONT direct RNA sequencing of Roche MS2 RNA library-prepped with poly(A)-tailing, poly(I)-tailing, or a custom adapter. **(D)** Per-base coverage for all three Nano-BID-Amp amplicons spanning the four U-to-C miscall sites of interest (Fig. 1C). Amplicon 1 spans genomic positions 346-1167, amplicon 2 spans genomic positions 1925-2533, and amplicon 3 spans genomic positions 2914-3569.

**Supplementary Figure 2.**
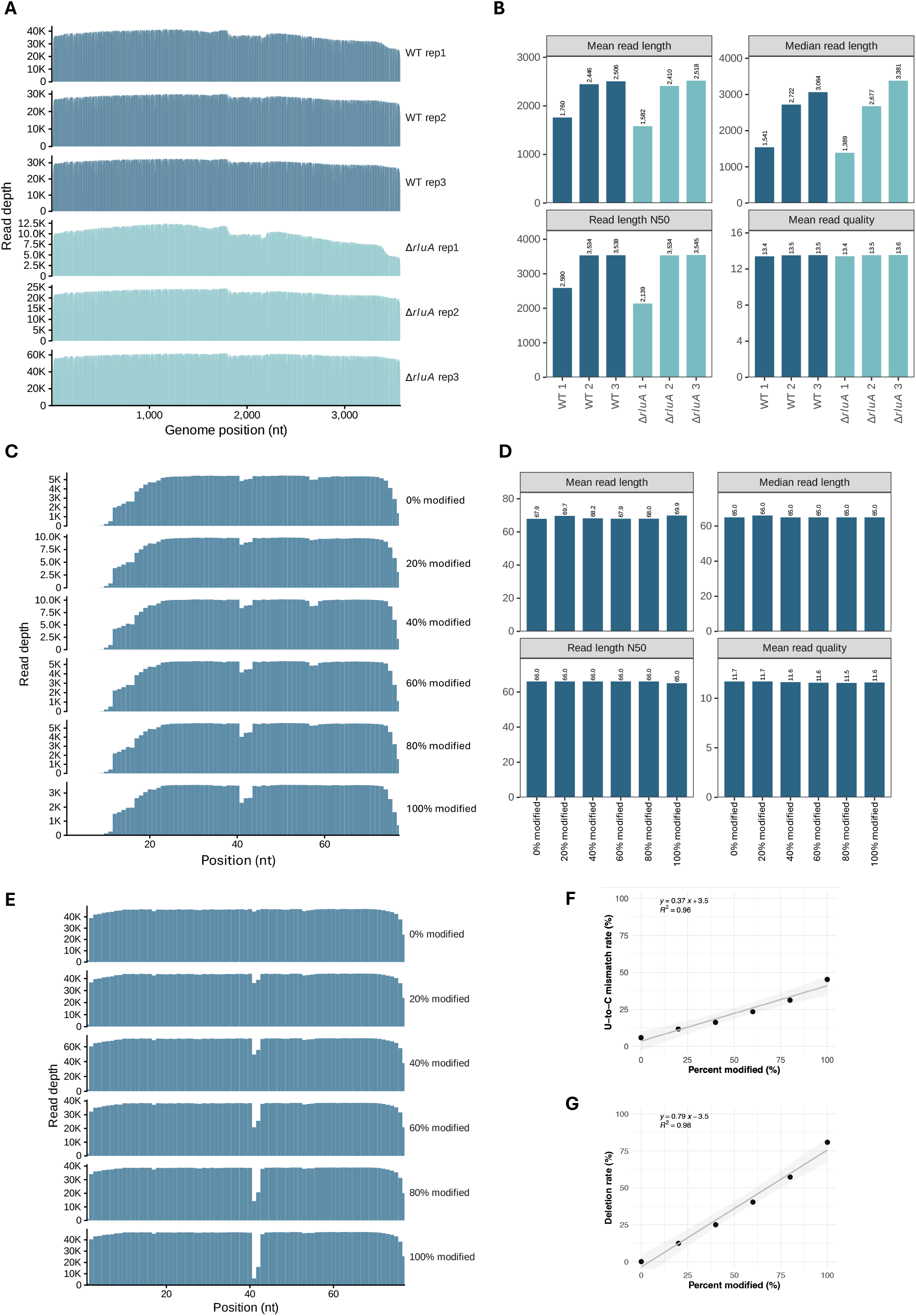
Sequencing coverage of MS2 from wild-type and ΔrluA host infections and of synthetic standards. **(A)** Per-base coverage of ONT direct RNA sequencing of MS2 from wild-type and Δ*rluA* hosts (three independent replicates). Replicate 2 was chosen as the representative data depicted in Fig. 2B and 2C. **(B)** Run statistics for ONT direct RNA sequencing of MS2 from wild-type and Δ*rluA* hosts (three independent replicates). **(C)** Per-base coverage of ONT direct RNA sequencing of physically mixed synthetic standards. **(D)** Run statistics for ONT direct RNA sequencing of synthetic standards. **(E)** Per-base coverage of Nano-BID-Amp sequencing of physically mixed synthetic standards. **(F)** Calibration curve for physical synthetic standard mix for ONT direct RNA sequencing. Gray shading represents 95% confidence intervals. **(G)** Calibration curve for physical synthetic standard mix for Nano-BID-Amp. Gray shading represents 95% confidence intervals.

**Supplementary Figure 3.**
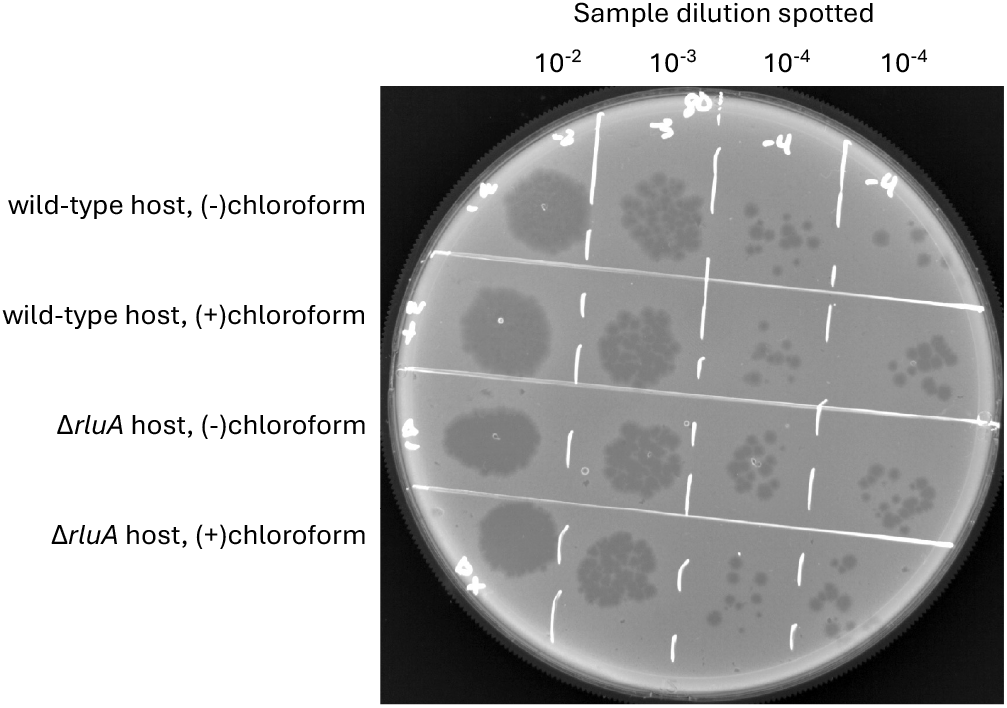
Enumerating PFU in one-step phage growth assay timepoints. Representative spotted plate for enumerating PFU in one-step phage growth assay timepoints. Spotting was performed in technical duplicates for each sample and each dilution.

**Supplementary Figure 4.**
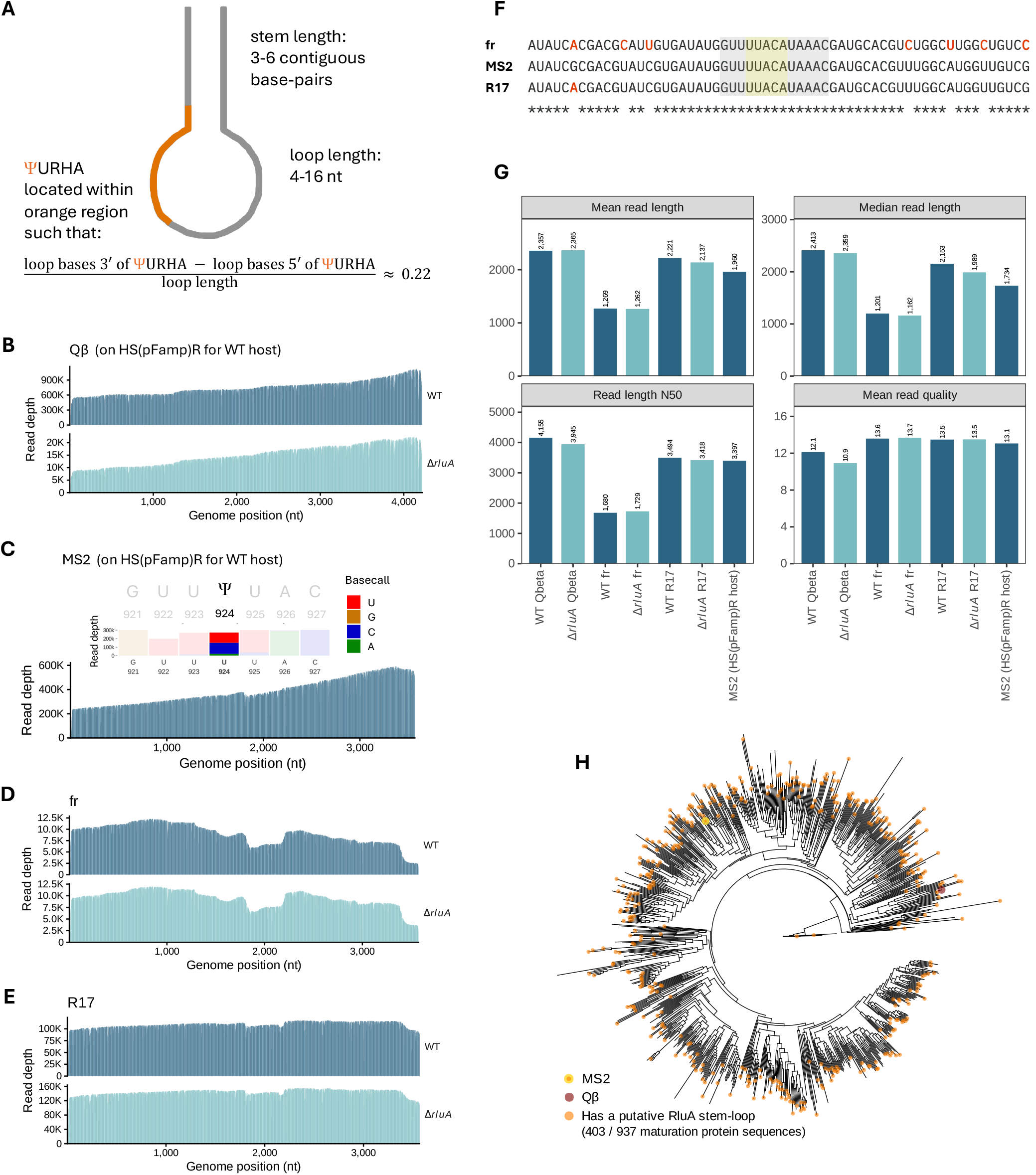
Features associated with RluA targets in other ssRNA phages. **(A)** Consensus RluA motif for the *in silico* search descriptor, derived from all known terminal-stem-loop RluA substrates with experimentally determined structures^75,76,43^. **(B)** Per-base coverage of ONT direct RNA sequencing of Qβ (strain from lab of Charlotte Knudsen) from wild-type and Δ*rluA* hosts (note different wild-type host: HS(pFamp)R (ATCC 700891)). **(C)** Top: ONT direct RNA sequencing basecall profile of genomic position 924 and flanking ±3 bases for MS2 from the same wild-type host as for Qβ, HS(pFamp)R (n = 1). Y-axis values indicate read depth. Bottom: Per-base coverage of ONT direct RNA sequencing of MS2 on host HS(pFamp)R. **(D)** Per-base coverage of ONT direct RNA sequencing of fr from wild-type and Δ*rluA* hosts. **(E)** Per-base coverage of ONT direct RNA sequencing of R17 from wild-type and Δ*rluA* hosts. **(F)** Alignment of MS2 Ψ924 region (genomic positions 898 to 957) with homologous R17 and fr regions. Bases in the R17 and fr sequences that differ from the MS2 sequence are in red. The RluA recognition sequence motif is highlighted in yellow and the RluA stem-loop is highlighted in gray. **(G)** Run statistics for ONT direct RNA sequencing of wild-type and Δ*rluA* Qβ, fr, and R17, as well as MS2 on the HS(pFamp)R host. **(H)** Maximum-likelihood tree depicting phylogeny of 937 *Leviviricetes* maturation protein sequences and distribution of *in silico* RluA target stem-loop hits. Orange tips indicate sequences with *in silico* hits. For reference, the yellow tip indicates MS2 phage and the maroon tip indicates Qβ phage.

## Materials & Methods

### Bacterial strains and culture conditions

Bacterial strains used in this study are listed in Table S1. The *ΔrluA E. coli* strain from the Keio collection^104^ was a kind gift from Rachel Porter and the lab of KC Huang. The Keio parent *E. coli* strain, BW25113, was a kind gift from Kelly Ann Lynch and the lab of Christine Jacobs-Wagner. *E. coli* HS(pFamp)R (ATCC 700891), a kind gift from Claire Anderson and the lab of Alexandra Boehm, was used as an F-plasmid donor.

The *ΔrluA F+ E. coli* strain and wild-type *F+ E. coli* strain (referred to simply as the wild-type host and the Δ*rluA* host throughout this study) were generated by conjugation of *E. coli* HS(pFamp)R with the *ΔrluA* and parent (BW25113) strains from the Keio collection^104^, using an adaptation of the Barrick lab’s conjugation protocol^105^. Overnight cultures of HS(pFamp)R (in LB with 100 µg/mL ampicillin, as the F plasmid encodes ampicillin resistance, and 100 µg/mL streptomycin, as the genome encodes streptomycin resistance) and recipient strains *ΔrluA* (in LB with 50 µg/mL kanamycin) and BW25113 (in LB) were grown to saturation. 1 mL of donor strain per 1 mL of each recipient strain was washed in PBS to remove residual antibiotic. The optical density was measured and each recipient strain was mixed with the donor strain at a 1:1 ratio. Each donor+recipient mixture was plated onto LB and incubated at 37 °C for three hours to ensure sufficient transfer of the F plasmid. Following conjugation, the mixture was scraped up, vortexed in PBS to release transconjugants, and serially diluted in additional PBS. Several of these ten-fold dilutions were plated on LB supplemented with 100 µg/mL ampicillin for the BW25113 recipient strain and with 100 µg/mL ampicillin and 50 µg/mL kanamycin for the *ΔrluA* recipient strain. Plates were incubated at 37 °C overnight. The following day, single transconjugant colonies were picked and restreaked onto both permissive plates (with ampicillin for the BW25113 recipient strain and with ampicillin and kanamycin for the *ΔrluA* recipient strain) and non-permissive plates with both ampicillin and streptomycin. Transconjugants that only grew on the permissive plates were verified with whole-genome sequencing to have two contigs—the chromosome and the F plasmid.

The resulting wild-type *F+* host strain and *ΔrluA F+* host strains were stored in 15% glycerol at-80 °C and streaked out freshly for each experimental replicate. For all subsequent phage-infections assays, both strains were grown on Tryptic Soy Agar (TSA; Fisher Scientific #DF0370-17-3) or in Tryptic Soy Broth (TSB; Fisher Scientific #DF0370-17-3) supplemented with 100 µg/mL ampicillin.

### Bacteriophage strains

Bacteriophage strains used in this study are listed in Table S1. Bacteriophage MS2 (ATCC 15597-B1) was a kind gift from Claire Anderson and the lab of Alexandra Boehm.

Bacteriophage Qβ and its genome sequence were kind gifts from Wadim Kapulkin and the lab of Charlotte Knudsen. Bacteriophage R17 (ATCC 25868-B1) and bacteriophage fr (ATCC 15767-B1) were obtained from ATCC.

The MS2 reference sequence, as well as the genomic coordinates shown in Fig. 1C, 2A, and S1A, were obtained from RefSeq (accession number NC_001417.2). We observed SNPs in our MS2 sequencing relative to this reference (Fig. S1A), and we note the often-imperfect sequence match between “real” isolates of RNA phages and their reference sequences^106^.

### Phage propagation (double-layer agar plaque assay)

Plaque assays were performed according to the double-layer agar method^80^. All phages were kept frozen in 15% glycerol at-80 °C and individual aliquots were thawed on ice for serial dilutions in SM buffer (G-Biosciences #786-491). Before transferring any volume of phage, the volume was pipetted up and down several times to reduce variability due to adsorption to plastic. All host strain growth media contained 100 µg/mL ampicillin. Molten soft agar was kept at ∼42-43 °C.

On the first day, host strains were streaked out onto tryptic soy agar (TSA) plates and incubated at 37 °C overnight. The following day, 5 mL TSB was inoculated with single colonies and grown overnight at 37 °C with shaking at 220 rpm. The following day, overnight cultures were back-diluted to OD_600_ = 0.05 in 5 mL TSB and allowed to grow at 37 °C with shaking at 220 rpm. Upon reaching OD_600_ ≈ 0.3 (early-to-mid-exponential phase), 200 µL of this culture was mixed with 300 µL of phage dilution. For the no-phage control (lawn) plates, 200 µL of the host was mixed with 300 µL of SM buffer. Adsorption of phage to host was allowed to proceed at room temperature for 10 minutes. Following this incubation, the phage-host mixture was added to 3.5 mL of soft TSA, mixed, and poured onto a prewarmed TSA plate. Plates were allowed to solidify and then incubated at 37 °C overnight (∼18 hours). Plates were imaged on a Bio-Rad GelDoc Go Imaging System using default “Fast Blast” settings with a white tray. Plaque-forming units (PFU) per mL were quantified as follows: (# of plaques / 300 µl) * 10^x (dilution factor)^ * 1000 µL/mL.

“Webbed” plates (plates with overlapping plaques) were selected for phage harvesting: three mL of ice-cold SM buffer (G-Biosciences #786-491) was added to the plate and allowed to incubate at 4 °C for three hours. The top soft agar layer was then gently broken up into the buffer, transferred with the buffer into a 15 mL conical tube, and centrifuged at 10,000 rpm at 4 °C for 30 minutes. The resulting supernatant was filtered twice through a 0.2 µm PES filter (Thermo Fisher) and stored in 15% glycerol at-80 °C.

The host strains for all phage-infection experiments were the wild-type *F+* and Δ*rluA F+* strains described above. The exception was Qβ, for which the “wild-type” F+ strain used at the time was *E. coli* HS(pFamp)R (ATCC 700891). We verified that growth on this wild-type *F+* host versus the wild-type *F+* host (BW25113) we used in our other experiments does not affect modification levels of MS2 (Fig. S4C; Table S3).

### RNA extraction

RNA was extracted from phage samples by adding TRIzol (ThermoFisher #15596018) at a 1:1 ratio, mixing thoroughly, and incubating for five minutes at room temperature. Chloroform was then added at 0.2 mL per 1 mL sample, mixed thoroughly, and incubated for three minutes at room temperature. Samples were centrifuged at 15,000 rpm for 15 minutes at 4 °C. The aqueous phase was transferred to a new tube on ice, to which ice-cold isopropanol was added at 0.5 mL per 1 mL sample and then mixed thoroughly. After incubation for at least 10 minutes on ice, samples were centrifuged at 15,000 rpm for 30 minutes at 4 °C. The supernatant was discarded, and the remaining RNA pellet was washed with ice-cold 70% ethanol and centrifuged at 15,000 rpm for 5 minutes at 4 °C. The supernatant was discarded, centrifuged briefly again, and any residual supernatant discarded. The remaining RNA pellet was resuspended in RNase-free water. Concentration was determined by Qubit RNA Broad Range Assay (ThermoFisher #Q10211).

3’-end-healing was performed by mixing ≤1 µg of extracted RNA with 4 µL low pH PNK buffer (0.5 M MES, pH 5.6; 50 mM MgCl_2_; 50 mM β-mercaptoethanol; 1.5 M NaCl), 2 µL 10 mM ATP, 1 µL T4 PNK (NEB #M0201S) in a total volume of 20 µL. Samples were incubated at 37 °C for 30 minutes and then cleaned up with Zymo RNA Clean & Concentrator-5 (Zymo Research #R1016).

Poly(A)-tailing was performed by assembling the following on ice in a total volume of 20 µL: ≤1 µg of extracted RNA, 2 µL 10X poly(A) polymerase buffer (NEB #B0276S), 2 µL 10 mM ATP, 1 µL poly(A) polymerase (NEB #M0276S). Samples were mixed, incubated at 37 °C for 30 seconds, and cleaned up with Zymo RNA Clean & Concentrator-5 (Zymo Research #R1016).

### MS2 synthetic standard design

Synthetic standards encompassing MS2 genomic positions 899 to 948 (50 bases) with or without a pseudouridine at genomic position 924 were ordered from Dharmacon, with a 3’ 10-nt poly(A) tail and a 5’ universal adapter (GACGCUCUUCCGAUCU).

### Oxford Nanopore Technologies (ONT) direct RNA sequencing

Initial direct RNA sequencing of MS2 genomic RNA depicted in Fig. 1C and S1A was performed with commercially obtained MS2 RNA (Roche #10165948001). The RNA was library-prepped using kit SQK-RNA004 with one of the following: poly(A)-tailing and the default oligo-d(T) adapter, poly(I)-tailing and a custom oligo-d(C) adapter^107^, or a custom adapter complementary to the 3’ end of the MS2 genomic RNA, and then sequenced on PromethION flow cells (Oxford Nanopore Technologies #FLO-PRO004RA).

Subsequent direct RNA sequencing of phages MS2, fr, and R17 from infection of wild-type and Δ*rluA* host strains was performed with RNA extracted from plaque assay plates as described above. This RNA was then 3’-end-healed and poly(A)-tailed. Libraries were prepared with kit SQK-RNA004 and barcoding kit SQK-DRB004.24 according to manufacturer guidelines and then sequenced on a PromethION flow cell (Oxford Nanopore Technologies #FLO-PRO004RA). Synthetic standard mixes of 100:0 to 0:100 ratios of unmodified:modified were included in the barcoding steps in order to generate calibration curves.

POD5 files from all direct RNA sequencing runs were basecalled with Dorado (version 0.9.1) using the high-accuracy model (rna004_130bps_hac@v3.0.1). Reads were mapped to the appropriate phage genome (MS2 RefSeq accession number NC_001417.2; R17 GenBank accession number EF108465.1; fr GenBank accession number GCA_002786405.1; Qβ reference genome provided by Wadim Kapulkin and the lab of Charlotte Knudsen) using minimap2 (`-ax map-ont`; version 2.27). Reads were then converted to BAM, sorted and indexed, and filtered for primary alignments (samtools version 1.9). ModDetect^58^, a custom in-house script, was used to identify all modified sites that manifest with base miscalls with a difference of ≥20% between wild-type reads and *ΔrluA* reads. For the three initial sequencing runs with commercially obtained MS2 RNA, a zero-error reference BAM file with 200 “perfect” reads was generated using the MS2 reference sequence (RefSeq accession number NC_001417.2), and positions with a reference U that displayed base miscalls (excluding indels) with a difference of ≥20% were outputted.

The synthetic standards were mapped using bowtie2 (version 2.3.4.3) with the following relaxed parameters to accommodate short nanopore reads: `--local --sensitive-local --ignore-quals --no-unal --mp 3,1--rdg 5,1 --rfg 5,1 --dpad 30-L 15`. Reads were then converted to BAM, sorted and indexed, and filtered for primary alignments (samtools version 1.9). A calibration curve for this physical mix was then plotted, from which a linear line of best fit equation was obtained. A computational mix was also generated by filtering 0%-modified and 100%-modified reads for read Q-scores ≥15 and then randomly subsampling both read sets at different modified:unmodified ratios. The total read count for all mix ratios was set to 90% of the read depth of the original 0%-and 100%-modified read pools. A calibration curve for these computationally mixed synthetic standards was obtained by generating the line of best fit. Modification stoichiometry estimates for MS2 phage samples were derived by substituting raw U-to-C miscall percentages into these equations. Stoichiometry estimates for phages fr and R17 were also obtained from the MS2-synthetic-standard calibration curves, given that the three phages share the same sequence identity in the pseudouridine site and flanking ±10 bases (Fig. S4F).

For Dorado modification calling, raw POD5 files were basecalled with Dorado (version 0.9.1) and the modification basecaller model rna004_130bps_sup@v5.1.0_pseU@v1. Unaligned reads were converted to fastq using samtools (`fastq-T *`; version 1.9) and aligned to the appropriate reference sequence with minimap2 (`-ax map-ont-y`; version 2.27). Modification estimates were obtained using modkit (`pileup` with `--mod=17802` (to specify pseudouridines), `--filter-threshold=0.9`, and `--mod-threshold=0.99`; version 0.4.0).

Basecall profile plots were visualized in R using samtools (version 1.9) mpileup.

### Illumina RNA-sequencing

Illumina cDNA sequencing data for MS2 RNA was obtained from a previous in-house sequencing run kindly shared by Drew Galls, which had included Roche MS2 RNA (Roche #10165948001) as a spike-in control. Briefly, the sequencing used a custom dual-end capture protocol that was inspired by several previously described approaches^108–110^: RNA was fragmented, ends were repaired, and adapters were ligated to both ends of the RNA. These adapters were used as handles for reverse transcription and PCR.

The RNA fragmentation reaction contained 250 ng RNA (MS2 RNA and other RNAs not described herein), 5 mM Na_2_CO_3_, 1 mM EDTA, and 45 mM NaHCO_3_ in a total volume of 10 μL. The reaction was incubated at 95 °C for 15 min and immediately transferred to ice. The fragmented RNA was purified with an Oligo Clean & Concentrator-5 kit (Zymo Research #D406) and eluted in 13 μL water. The 3’-end-clean-up reaction was then performed as described above. RNA was purified with an Oligo Clean & Concentrator-5 kit (Zymo Research #D406) and eluted 6 μL IDTE pH 7.5 (IDT #11-05-01-15).

The 3’ adapter was prepared as previously described^110^. The 3’ ligation reaction contained 0.6 μM 3’ adapter (6 pmol AF-DG-214; Table S2), 20,000 U/mL T4 RNA Ligase 2 truncated KQ (NEB #M0373; 200 U), 20% PEG-8000, and T4 RNA Ligase Buffer (NEB #B0216) in a total volume of 10 μL. 40 ng RNA and adapter were combined in a 4 μL volume. The mixture was heated to 70 °C for 2 min and cooled on ice, and 6 μL of the ligase, PEG-8000, and ligase buffer mixture was added. The reaction was incubated at 30 °C for 1 hour. Post-ligation, adapters were degraded in a reaction containing 500 U/mL 5’ deadenylase (NEB #M0331; 50 U), 300 U/mL RecJf (NEB #M0264; 30 U), the ligation reaction, and 1x NEBuffer 2 (NEB #B7002) in a total volume of 100 μL. The reaction was incubated at 30 °C for 15 min and then at 37 °C for 15 min. Ligated RNA was purified with an Oligo Clean & Concentrator-5 kit (Zymo Research #D406) using the protocol for RNAs >80 nt and then eluted in 6 μL T4 RNA Ligase Buffer (NEB #B0216).

For the 5’ ligation reaction, the following were combined in a total volume of 12 µL: 1.67 μM 5’ adapter (20 pmol AF-DG-220; Table S2), 833 U/mL T4 RNA Ligase 1 (NEB #M0204; 10 U), 1.25 mM ATP, 19.4% PEG-8000, and T4 RNA Ligase Buffer (NEB #B0216). The 5’ adapter was first heated to 72 °C for 2 min and cooled on ice, and then the enzyme, ATP, and PEG were added. 6 μL of this mixture was then added to the 6 μL of RNA in T4 RNA Ligase Buffer (NEB #B0216), and this resulting 5’ ligation reaction was incubated at 25 °C for 1 hour.

For the RT reaction, the following were combined in a total volume of 25 µL: 0.32 μM RT primer (8 pmol AF-DG-3; Table S2), 500 μM dNTPs, 8 mM DTT, 16,000 U/mL ProtoScript II (NEB #M0368; 400 U), 0.8x ProtoScript II Buffer (NEB #B0368), and the entire volume of the ligation reaction. First, 2 μL RT primer was added to the 12 µL ligation reaction, the reaction was heated to 72 °C and cooled on ice, the mixture of buffer, dNTPs, DTT, and enzyme was added, and the reaction was incubated at 42 °C for 1 hour.

The PCR reaction contained 0.24 μM of each primer (24 pmol each), 1x NEBNext High-Fidelity 2X PCR Master Mix (NEB #M0541), and the entire volume of the RT reaction in a total volume of 100 μL. The PCR was separated into four 25 μL reactions to minimize jackpotting. Cycling conditions were: 12 cycles of 98 °C for 10 sec, 60 °C for 30 sec, and 72 °C for 15 sec; initial denaturation at 98 °C for 30 sec; final extension at 72 °C for 15 min. Amplified DNA was purified with a DNA Clean & Concentrator-5 kit (Zymo Research #D4014) and eluted in 15 μL IDTE pH 8.0 (IDT 11-05-01-13). Gel purification and library quantification were as previously described^110^, except the gel was 3% agarose and fragments of ∼200-500 nt were excised. Libraries were sequenced by Azenta. Subsequently, an index was built from the MS2 reference sequence (RefSeq accession number NC_001417.2) with bowtie2 (version 2.3.4.3) and fastq files were aligned (`--very-sensitive-local`).

### Nano-BID-Amp

The Nano-BID-Amp protocol was adapted from previously published protocols^64,58^, with the addition of synthetic standards.

Twenty ng of wild-type and Δ*rluA* MS2 RNA in a total volume of 8.5 µl nuclease-free water each were transferred to PCR strip tubes. A bisulfite mix resulting in 2.4 M Na_2_SO_3_ and 0.36 M NaHSO_3_ was resuspended in 1 mL of nuclease free water with vigorous vortexing. Immediately after bisulfite resuspension, 45 μl of bisulfite mix was added to RNA samples previously equilibrated to room temperature. Samples were mixed well and incubated at 70 °C for 3 hours. The Zymo RNA Clean and Concentrator-5 kit (Zymo Research #R1016) was used to purify RNA by mixing the samples with 75 μl water, 270 μl RNA Binding Buffer (Zymo Research #R1013-2-100), and 400 μl 100% ethanol; loading onto columns attached to a vacuum manifold via sterile adapters; and vacuuming through. Samples were then washed once with 200 μl RNA Wash Buffer (Zymo Research #R1003-3-48) and the wash buffer was discarded by vacuum manifold. 200 μl of RNA Desulphonation Buffer (Zymo Research #R5001-3-40) was added directly to the columns and samples were incubated at room temperature for 75 minutes. Desulphonation buffer was discarded via vacuum manifold and samples were washed twice with RNA Wash Buffer (700 μl for first wash, 400 μl for second wash). Samples were spun for 1 minute at 16,000 x g to remove any residual buffer not removed by the vacuum manifold, eluted in 11 μl nuclease free water, and immediately proceeded to reverse transcription.

For reverse transcription, 10.5 µl of the bisulfite-treated RNA and 10.5 µl of untreated input control (20 ng of both wild-type and Δ*rluA* MS2 RNA) were mixed with 1 µl of 2 µM template-specific primers with a 10-mer UMI and ONT sequencing adapters. Samples were incubated at 65 °C for 2 minutes, and then held at 50 °C while the reverse transcription master mix was added and mixed (4 μl 5X SuperScript IV Buffer (Invitrogen #18090050), 2 μl 10 mM dNTPs, 1 μl 100 mM DTT, 0.5 μl RNasin Plus (Promega #N2611), and 1 μl SuperScript IV (Thermo Fisher #18090200)). Samples were incubated at 50 °C for 1 hour. Following incubation, each sample was mixed with 2 µl of 4M NaOH and incubated at 95 °C for 3 minutes to degrade the RNA, and then immediately proceeded to clean-up with a 1.5X volume of RNAClean XP beads (Beckman Coulter #A63987) for phage samples and 2X volume for synthetic standards. Bead clean-up was repeated to fully remove UMI-containing primers.

4 µl of purified cDNA was added to a final PCR volume of 20 µl (4 µL 5X Phusion HF buffer (New England Biolabs #B0518S), 4 µL 10 mM dNTPs, 1 µl of each 10 µM primer (with ONT adapter sequences), and 2 µl Phusion polymerase (New England Biolabs #M0530L)), and proceeded through ONT-tailing PCR (18 cycles for input samples, 20 cycles for bisulfite-treated samples). Resulting PCR products were verified with a TapeStation D1000 ScreenTape (Agilent #5067–5582) before clean-up with 1.8X volume AMPure XP beads (Beckman Coulter #A63881) and elution in 20 µl of nuclease-free water. Samples were then barcoded with the Nanopore cDNA Ligation kit (Oxford Nanopore Technologies #SQK-LSK114) and the PCR Barcoding Expansion kit (Oxford Nanopore Technologies #EXP-PBC096) with 13 cycles of barcoding PCR. These barcoded samples were pooled and sequenced on a PromethION (Oxford Nanopore Technologies #FLO-PRO114M) flow cell.

Synthetic standard mixes of 100:0 to 0:100 ratios of unmodified:modified were included in this pipeline in order to generate a calibration curve.

Post-sequencing, reads were processed from fastq files by adapting a custom in-house script^58^. Reads were filtered and trimmed with cutadapt (`-m 125-O15 --discard-untrimmed` for phage reads; `-m 20-O15 --discard-untrimmed` for synthetic standards; version 5.2). The orientation of all reads was converted to sense with seqtk (version 1.5) before extracting UMIs with umi_tools (`extract`; version 1.1.6). Phage reads were mapped with minimap2 (`-ax map-ont`; version 2.31). Synthetic standard reads were filtered for read Q-scores ≥15 and mapped with bowtie2 (version 2.5.5) with the following relaxed parameters to accommodate short nanopore reads: `--local --sensitive-local -- ignore-quals --no-unal --mp 3,1 --rdg 5,1 --rfg 5,1 --dpad 30-L 15`. Reads were then converted to BAM, sorted and indexed with samtools (version 1.24), filtered for primary alignments only with a MAPQ score ≥30, and deduplicated with umi_tools (`dedup – method directional`; version 1.1.6). The deletion fraction value for the input (untreated) was subtracted from the deletion fraction value for its corresponding bisulfite-treated sample. A calibration curve for this physical mix was then plotted, from which a linear line of best fit equation was obtained.

The computational synthetic standard mix was generated by filtering both untreated and bisulfite-treated 0%-modified and 100%-modified reads for read Q-scores ≥15 and then randomly subsampling all read sets at different modified:unmodified ratios. The total read count for all mix ratios was set to 90% of the read depth of the original 0%-and 100%-modified read pools. The resulting untreated values were subtracted from corresponding bisulfite-treated values. A calibration curve for these computationally mixed synthetic standards was obtained by generating the line of best fit. Modification stoichiometry estimates were derived by substituting raw deletion rates into this equation.

Basecall profile plots were visualized in R using samtools (version 1.9) mpileup.

### Adsorption efficiency assay

5 ml cultures inoculated with single colonies of host strains were grown at 37 °C shaking at 220 rpm overnight. The subsequent day, 20 ul of each culture was back-diluted into 5 ml TSB (supplemented with 100 µg/ml ampicillin, 5 mM CaCl_2_, 10 mM MgCl_2_) and grown at 37 °C shaking at 220 rpm until reaching OD_600_ = 0.3-0.4 (early-to-mid-log phase). MS2 phage was mixed with ∼2.5 x 10^8^ cells in 1 mL at an MOI of 0.01 and allowed to adsorb at 37 °C for 10 minutes. Samples were then spun for five minutes at 13,000 x g. The PFU of the supernatant, which contains unadsorbed phage, was assessed by spotting assay of ten-fold serial dilutions. Unadsorbed phage fraction was calculated relative to the input of 2.5 x 10^6^ PFU.

### One-step phage growth assay

5 ml cultures inoculated with single colonies of host strains were grown at 37 °C shaking at 220 rpm overnight. The subsequent day, 20 µl of each culture was back-diluted into 5 ml TSB (supplemented with 100 µg/ml ampicillin, 5 mM CaCl_2_, 10 mM MgCl_2_) and grown at 37 °C shaking at 220 rpm until reaching OD_600_ = 0.3-0.4 (early-to-mid-log phase). MS2 phage was mixed with ∼2.5 x 10^8^ cells in 1 mL at an MOI of 0.01 and allowed to adsorb at 37 °C for 10 minutes. Following adsorption, samples were washed twice with fresh media to remove unadsorbed phage, diluted 1000-fold into 50 mL fresh media, and incubated at 37 °C with shaking at 220 rpm. Two 500 µl samples of both wild-type and *ΔrluA* host infection cultures were taken every 5-10 minutes. 10 µl chloroform was added to one sample per host strain. All samples were then vortexed briefly, incubated at room temperature for one minute, spun down for three minutes at 13,000 x g, and placed on ice until plating for PFU enumeration immediately after completion of sample collection. PFUs for all timepoints were enumerated either by spotting assay with technical duplicates (5 µl sample dilution spotted in duplicate onto 3 ml soft agar overlay mixed with 100 µl early-to-mid-log phase wild-type *F+* host (OD_600_ = 0.3-0.4)) or by full-plate plaque assay as described above.

To estimate burst size, PFUs for each host strain at post-adsorption timepoints 110 and 120 minutes (where the curve was observed to plateau) were averaged and then normalized by their respective averaged infective centers determined via adsorption efficiency assay. For estimations of both the minimum latent period and the eclipse period that do not rely on visual approximation, the threshold was set as 10^2^ PFU/mL and the time of crossing this threshold was interpolated from sampled timepoints. For all strains with or without chloroform-induced lysis, a line was drawn between the first timepoint above that threshold and the previous timepoint and was used to calculate the intercept at 10^2^ PFU/mL. The-chloroform timepoints were used to estimate each strain’s latent period and the +chloroform timepoints were used to estimate each strain’s eclipse period.

### Growth curve

5 ml cultures inoculated with single colonies of host strains were grown at 37 °C shaking at 220 rpm overnight. The subsequent day, 20 µl of each culture was back-diluted into 5 ml TSB (supplemented with 100 µg/ml ampicillin, 5 mM CaCl_2_, 10 mM MgCl_2_) and grown at 37 °C shaking at 220 rpm until reaching OD_600_ = 0.3-0.4 (early-to-mid-exponential phase). These log-phase cultures were back-diluted to a starting OD_600_ of 0.02 into 500 µl TSB (100 µg/ml ampicillin), with two technical replicates in a 48-well plate (Corning #351178). Growth curves were allowed to proceed for 18 hours in a Tecan Infinite 200 PRO with the following settings: interval every 10 minutes: orbital shaking (420 seconds at amplitude 3), wait (5 seconds), absorbance reading (600 nm; 25 flashes, 50 ms settle time). All readings were normalized to blank wells. Log-phase doubling times were estimated from a linear line of best fit generated from datapoints between 0.5 and 2.5 hours.

### Homology search for unannotated pseudouridine-synthase-like domain

To search for unannotated pseudouridine-synthase-like domains in all four MS2 coding sequences, the following programs were used with default settings: hmmscan (search for sequence homologs; HMMER version 3.4), InterProScan (domain prediction; version 5.78), CD-Search (search for conserved domains; version 3.21), HHPred (search for more remote homology). No search yielded hits with percent identities >50% and E-values < 0.05 that were not to the expected maturation protein, coat protein, lysis protein, or RNA-dependent RNA polymerase.

### ssRNA coliphage sequence alignment and phylogenetic tree construction

Alignment of “classic” ssRNA coliphage genome sequences and calculation of percent identity (--percent-id) was performed with Clustal Omega (version 2.1) with default settings.

A tree illustrating sequence identity relationships between these genomes was generated by multiple sequence alignment of genome sequences with MAFFT (version 7.525) default settings, converted to PHYLIP format with easel (version 0.49), and then processed with FastME (version 2.1.6.3) with default settings, which computes pairwise distance and then builds an unrooted tree around ‘minimum evolution’ constraints. The tree was visualized with ITOL Interactive Tree of Life^111^ (version 7.6), without branch lengths.

Alignment of maturation protein sequences was performed by translating the nucleotide sequences to amino acids with seqkit (translate --trim; version 0.10.0), aligning with MAFFT (--auto; version 7.525), and trimming with TrimAl (-gt 0.3; version 1.5.0).

### RluA stem-loop definition and search

The MS2 Ψ924 stem-loop structural prediction was generated with ViennaRNA (version 2.6.3) RNAfold using windows of ±50, ±100, and ±150 flanking bases. These predicted structures were identical.

For the broader *in silico* search: All known terminal stem-loop substrates of RluA with experimentally determined secondary structures were included when determining shared features^43,75,76^ (23S rRNA:ψ746, tRNA-Phe:ψ32, tRNA-Cys:ψ32, tRNA-LeuX:ψ32, tRNA-LeuZ:ψ32, *pepD:*ψ1024, *pyrC:*ψ75, *rpsA:*ψ460, *rpsA:*ψ176, *hpf:*ψ58, *fadL:*ψ538, *nuoG:*ψ2356, *fur:*ψ57, *ygiC:*ψ133, *sslE:*ψ1909, *rpsA:*ψ880, *iscS:*ψ335, *glf:*ψ592, *gtrS:*ψ156, *slyA:*fi342). The mean and median ratio of *(loop bases 3’ of UURHA - loop bases 5’ of UURHA) / (loop length)* was ∼0.22. We defined the RluA substrate descriptor as follows for our search: UURHA is situated within the loop such that the first U is located in the closing base-pair when the loop is 4-5 nt, 1 base into the loop when the loop is 6-7 nt, 2 bases into the loop when the loop is 8-10 nt, 2-4 bases into the loop when the loop is 11-14 nt, and 2-5 bases into the loop when the loop is 15-16 nt. For a more conservative search, we excluded outliers using Tukey’s fences: loop length was limited to values within 1.5 x interquartile range (4-16 nt) and qualifying stem length was limited to values within 1.5 x interquartile range (3-6 contiguous nt) (Fig. S4A, Table S8).

All *Leviviricetes* maturation protein gene sequences were retrieved from NCBI Virus as follows:

`esearch-db nuccore-query ‘txid2842243[Organism:exp] AND srcdb_refseq[PROP]’| efetch-format fasta_cds_na`

followed by:

`awk ‘/^>/{h=tolower($0); keep = (h ∼ /\[protein=[^]]*(maturation|assembly|attachment|A|A2)/) || (h ∼ /\[protein=([^]]*[^a-z0-9])?a2?[-]*protein/)} keep’`

(The maturation protein is also referred to as the maturation protease, attachment protein, assembly protein, A protein, or A2 protein.)

These sequences were searched for matches to the motif criteria using a custom R script. In short, sequences were computationally folded with RNALfold (version 2.7.2) using a folding window of 50 nt. All outputted structures were scanned for UURHA regex matches. If the UURHA was located within the correct structural context as defined above, the stem-loop was outputted as a putative RluA target hit. In the instance that the same UURHA motif site appeared in multiple overlapping predicted structures, only the structure with the lowest predicted free energy was outputted as the hit.

### Leviviricetes maturation protein gene phylogenetic tree construction

All *Leviviricetes* maturation protein gene sequences (nucleotide sequences) retrieved from NCBI Virus above were translated to amino acids with seqkit (`translate --trim`; version 0.10.0), aligned with MAFFT (`--auto`; version 7.525), and trimmed with TrimAl (`-gt 0.3`; version 1.5.0), which removed alignment columns in which over 70% of sequences contained a gap and thereby reduced the total columns from 4,682 to 447. A maximum-likelihood tree was inferred with IQ-TREE 2 (version 2.3.6), using ModelFinder (`-m MFP`) with the default Bayesian Information Criterion to determine the best-fit model. The resulting model (Q.pfam+F+R10) was used to build a maximum-likelihood tree with concurrent ultrafast bootstrap approximation (1000 replicates; `-bb 1000`). The median ultrafast bootstrap support value across all nodes was 99, with 619 of 934 internal nodes at ≥95. Tree is unrooted.

## Data representation

All plots were generated using R (version 4.4.3) unless otherwise indicated, with the following packages: ggplot2 (version 4.0.3), ggh4x (version 0.3.1), ggpubr (version 1.0.0), dplyr (version 2.6.0), tidyr (version 1.3.2), ggsignif (version 0.6.4), broom (version 1.0.13), ggseqlogo (version 0.2.2), Biostrings (version 2.74.0), Rsamtools (version 2.22.0), ggcoverage (version 0.7.1), ggtree (version 3.14.0), scales (version 1.4.0), ape (version 5.8.1). RNA secondary structure diagrams were generated with ViennaRNA RNAplot (version 2.4.7).

## Code and data availability

Code associated with this study can be found here: https://github.com/janieshk/phage-RNA-mods/

Raw sequencing data associated with this study can be found here: https://purl.stanford.edu/md919gh6774

## Declaration of generative AI in coding

The Stanford AI Playground (Anthropic Claude Opus 4.8) and Stanford Claude for Education (Opus 5) were used to assist with coding. All suggestions from AI were reviewed for accuracy and then adopted, adapted, or discarded.

## Acknowledgments

We thank Claire Anderson and the lab of Alexandra Boehm for gifting MS2 phage ATCC 15597-B1 and host strains ATCC 700891 and ATCC 15597; Wadim Kapulkin and the lab of Charlotte Knudsen for gifting their strain of Qβ phage and its genome sequence; Rachel Porter and the lab of KC Huang for gifting the Δ*rluA* Keio strain; Kelly Ann Lynch and the lab of Christine Jacobs-Wagner for gifting the Keio parent strain, BW25113. We thank Drew Galls for sharing MS2 Illumina sequencing data and protocol and for pointing us towards the 1970s MS2 sequencing papers. We thank past and present members of the Martinez and Fire labs—Karen Artiles, Colette Benko, Usman Enam, Drew Galls, Zoya Gauhar, Emily Greenwald, Orkan Ilbay, Ronit Jain, Dae-Eun Jeong, Haim Krupkin, Matt McCoy, Matias Montes, Nico Robalin, Becca Rodell, Christina Rouhotas, Hossein Shenasa, Wendy Trieu, Kailey Worner, and Vanya Zheludev—for valuable discussion and critical feedback. We also thank Devaki Bhaya, Mohamed Donia, Wadim Kapulkin, Charlotte Knudsen, Roberto Kolter, Jin Billy Li, Sharon Long, Lavisha Parab, Amanda Shelton for their valuable discussion and feedback on the manuscript.

N.M.M. is a Biohub, San Francisco, Investigator. Funding was provided by a Rita Allen Foundation Award and a Packard Fellowship for Science and Engineering from the David and Lucile Packard Foundation to N.M.M; by NIGMS-NIH (R35GM130366) to A.Z.F; and by an NSF Graduate Research Fellowship (DGE-1656518) and Ruth L. Kirschstein NRSA Individual Fellowship (NIAID-NIH: F31AI197524) to J.S.K.

The content is solely the responsibility of the authors and does not necessarily represent the official views of the National Institutes of Health and of the National Science Foundation.

## Author Contributions

J.S.K., A.Z.F., and N.M.M. conceived the study and designed experiments. J.S.K. performed ONT direct RNA sequencing, Nano-BID-Amp, phage propagation, strain construction, plaque assays, one-step phage growth assays, adsorption assays, *in silico* motif search, data analyses, and data visualization. H.S. performed ONT direct RNA sequencing and Nano-BID-Amp with synthetic standards. J.S.K. wrote the manuscript original draft, and J.S.K., H.S., A.Z.F., and N.M.M. reviewed and edited the manuscript. A.Z.F. and N.M.M. supervised the study. All authors approved the final manuscript.

## References

1. Haegeman, G. & Fiers, W. Studies on the Bacteriophage MS2: An Internal Nucleotide Fragment Resembling Some Ribosomal Binding Sites. Eur. J. Biochem. 36, 135–143 (1973).

2. Fiers, W. et al. A-Protein gene of bacteriophage MS2. Nature 256, 273–278 (1975).

3. Fiers, W. et al. Complete nucleotide sequence of bacteriophage MS2 RNA: primary and secondary structure of the replicase gene. Nature 260, 500–507 (1976).

4. Callanan, J. et al. Expansion of known ssRNA phage genomes: From tens to over a thousand. Sci. Adv. 6, eaay5981 (2020).

5. Callanan, J. et al. Leviviricetes: expanding and restructuring the taxonomy of bacteria-infecting single-stranded RNA viruses. *Microb*. Genomics 7, (2021).

6. Zayed, A. A. et al. Cryptic and abundant marine viruses at the evolutionary origins of Earth’s RNA virome. Science 376, 156–162 (2022).

7. Neri, U. et al. Expansion of the global RNA virome reveals diverse clades of bacteriophages. Cell 185, 4023–4037.e18 (2022).

8. Turner, D. et al. Abolishment of morphology-based taxa and change to binomial species names: 2022 taxonomy update of the ICTV bacterial viruses subcommittee. Arch. Virol. 168, 74 (2023).

9. Robertson, H. D. & Lodish, H. F. Messenger Characteristics of Nascent Bacteriophage RNA. Proc. Natl. Acad. Sci. 67, 710–716 (1970).

10. Schmidt, B. F., Berkhout, B., Overbeek, G. P., Van Strien, A. & Van Duin, J. Determination of the RNA secondary structure that regulates lysis gene expression in bacteriophage MS2. J. Mol. Biol. 195, 505–516 (1987).

11. Peabody, D. S. Translational repression by bacteriophage MS2 coat protein expressed from a plasmid. A system for genetic analysis of a protein-RNA interaction. J. Biol. Chem. 265, 5684–5689 (1990).

12. Poot, R. A., Tsareva, N. V., Boni, I. V. & Van Duin, J. RNA folding kinetics regulates translation of phage MS2 maturation gene. Proc. Natl. Acad. Sci. 94, 10110–10115 (1997).

13. Licis, N. Long-range translational coupling in single-stranded RNA bacteriophages: an evolutionary analysis. Nucleic Acids Res. 26, 3242–3246 (1998).

14. Gytz, H. et al. Structural basis for RNA-genome recognition during bacteriophage Qβ replication. Nucleic Acids Res. 43, 10893–10906 (2015).

15. Tars, K. ssRNA Phages: Life Cycle, Structure and Applications. in Biocommunication of Phages (ed. Witzany, G.) 261–292 (Springer International Publishing, Cham, 2020). doi:10.1007/978-3-030-45885-0_13.

16. Wagner, A., Weise, L. I. & Mutschler, H. In vitro characterisation of the MS2 RNA polymerase complex reveals host factors that modulate emesviral replicase activity. *Commun*. Biol. 5, 264 (2022).

17. Valegård, K., Liljas, L., Fridborg, K. & Unge, T. The three-dimensional structure of the bacterial virus MS2. Nature 345, 36–41 (1990).

18. Koning, R. I. et al. Asymmetric cryo-EM reconstruction of phage MS2 reveals genome structure in situ. Nat. Commun. 7, 12524 (2016).

19. Karikó, K., Buckstein, M., Ni, H. & Weissman, D. Suppression of RNA Recognition by Toll-like Receptors: The Impact of Nucleoside Modification and the Evolutionary Origin of RNA. Immunity 23, 165–175 (2005).

20. Roundtree, I. A., Evans, M. E., Pan, T. & He, C. Dynamic RNA Modifications in Gene Expression Regulation. Cell 169, 1187–1200 (2017).

21. Boccaletto, P. et al. MODOMICS: a database of RNA modification pathways. 2017 update. Nucleic Acids Res. 46, D303–D307 (2018).

22. Gilbert, W. V. & Nachtergaele, S. mRNA Regulation by RNA Modifications. Annu. Rev. Biochem. 92, 175–198 (2023).

23. Davis, D. R., Veltri, C. A. & Nielsen, L. An RNA Model System for Investigation of Pseudouridine Stabilization of the Codon-Anticodon Interaction in tRNA^Lys^, tRNA^His^ and tRNA^Tyr^. J. Biomol. Struct. Dyn. 15, 1121–1132 (1998).

24. Sipa, K. et al. Effect of base modifications on structure, thermodynamic stability, and gene silencing activity of short interfering RNA. RNA 13, 1301–1316 (2007).

25. Kierzek, E. et al. The contribution of pseudouridine to stabilities and structure of RNAs. Nucleic Acids Res. 42, 3492–3501 (2014).

26. Chen, C., Zhao, X., Kierzek, R. & Yu, Y.-T. A Flexible RNA Backbone within the Polypyrimidine Tract Is Required for U2AF^65^ Binding and Pre-mRNA Splicing *In Vivo*. Mol. Cell. Biol. 30, 4108–4119 (2010).

27. Banerjee, D. et al. C-Nucleosides Stabilize RNA by Reducing Nucleophilicity at 2′-OH. ACS Cent. Sci. 11, 2400–2409 (2025).

28. Islam, Md. S., Bandyra, K. J., Chao, Y., Vogel, J. & Luisi, B. F. Impact of pseudouridylation, substrate fold, and degradosome organization on the endonuclease activity of RNase E. RNA 27, 1339–1352 (2021).

29. Gutierrez, C. S. et al. Pseudouridine Residues as Substrates for Serum Ribonucleases. RNA rna. 080404.125 (2025) doi:10.1261/rna.080404.125.

30. Anderson, B. R. et al. Nucleoside modifications in RNA limit activation of 2’-5’-oligoadenylate synthetase and increase resistance to cleavage by RNase L. Nucleic Acids Res. 39, 9329–9338 (2011).

31. Durbin, A. F., Wang, C., Marcotrigiano, J. & Gehrke, L. RNAs Containing Modified Nucleotides Fail To Trigger RIG-I Conformational Changes for Innate Immune Signaling. mBio 7, e00833–16 (2016).

32. Vaidyanathan, P. P., AlSadhan, I., Merriman, D. K., Al-Hashimi, H. M. & Herschlag, D. Pseudouridine and *N*^6^-methyladenosine modifications weaken PUF protein/RNA interactions. RNA 23, 611–618 (2017).

33. Levi, O. & Arava, Y. S. Pseudouridine-mediated translation control of mRNA by methionine aminoacyl tRNA synthetase. Nucleic Acids Res. 49, 432–443 (2021).

34. Hoernes, T. P. et al. Nucleotide modifications within bacterial messenger RNAs regulate their translation and are able to rewire the genetic code. Nucleic Acids Res. 44, 852–862 (2016).

35. Anderson, B. R. et al. Incorporation of pseudouridine into mRNA enhances translation by diminishing PKR activation. Nucleic Acids Res. 38, 5884–5892 (2010).

36. Pozhydaieva, N., Wolfram-Schauerte, M., Keuthen, H. & Höfer, K. The enigmatic epitranscriptome of bacteriophages: putative RNA modifications in viral infections. Curr. Opin. Microbiol. 77, 102417 (2024).

37. Guegler, C. K. & Laub, M. T. Shutoff of host transcription triggers a toxin-antitoxin system to cleave phage RNA and abort infection. Mol. Cell 81, 2361–2373.e9 (2021).

38. Azam, A. H. et al. Evasion of antiviral bacterial immunity by phage tRNAs. Nat. Commun. 15, 9586 (2024).

39. Nikolic, N., Pleška, M., Bergmiller, T., & Călin C Guet. A bacterial toxin–antitoxin system as a native defence element against RNA phages. Biol. Lett. 21, (2025).

40. Drobysheva, A., Gomariz, M. V., Ahmad, S., Reichardt, S. & Hör, J. ApeA cleaves genomic RNA to defend against RNA phage infection.

41. Chihara, K. et al. Phage homing endonuclease amplifies anti-defense genes to evade bacterial immunity. Nat. Commun. 17, 3468 (2026).

42. Deng, X. et al. Widespread occurrence of *N* ^6^-methyladenosine in bacterial mRNA. Nucleic Acids Res. 43, 6557–6567 (2015).

43. Schaening-Burgos, C., LeBlanc, H., Fagre, C., Li, G.-W. & Gilbert, W. V. RluA is the major mRNA pseudouridine synthase in Escherichia coli. PLOS Genet. 20, e1011100 (2024).

44. Sharma, S. et al. Quantitative mapping of pseudouridines in bacterial RNA. Nat. Commun. 17, 3242 (2026).

45. Petrov, D. P., Kaiser, S., Kaiser, S. & Jung, K. Opportunities and Challenges to Profile mRNA Modifications in Escherichia coli. ChemBioChem 23, e202200270 (2022).

46. Xu, L. et al. Quantitative RNA pseudouridine landscape reveals dynamic modification patterns and evolutionary conservation across bacterial species. eLife 14, RP107545 (2026).

47. Fasnacht, M. et al. Dynamic 23S rRNA modification ho5C2501 benefits *Escherichia coli* under oxidative stress. Nucleic Acids Res. 50, 473–489 (2022).

48. Fruci, M. et al. Structural Analysis of 23S rRNA Methylating Enzyme Cfr Reveals RNA-Binding Determinants for Methylation Regiospecificity and Antibiotic Resistance. ACS Infect. Dis. 12, 653–664 (2026).

49. Giessing, A. M. B. et al. Identification of 8-methyladenosine as the modification catalyzed by the radical SAM methyltransferase Cfr that confers antibiotic resistance in bacteria. RNA 15, 327–336 (2009).

50. Keffer-Wilkes, L. C., Veerareddygari, G. R. & Kothe, U. RNA modification enzyme TruB is a tRNA chaperone. Proc. Natl. Acad. Sci. 113, 14306–14311 (2016).

51. Babosan, A. et al. Nonessential tRNA and rRNA modifications impact the bacterial response to sub-MIC antibiotic stress. Gene Funct. (2022).

52. Tsai, K. et al. Directed evolution of the rRNA methylating enzyme Cfr reveals molecular basis of antibiotic resistance. eLife 11, e70017 (2022).

53. Saikia, B., Riquelme-Barrios, S., Carell, T., Brameyer, S. & Jung, K. Depletion of m^6^ A-RNA in *Escherichia coli* reduces the infectious potential of T5 bacteriophage. Microbiol. Spectr. 12, e01124–24 (2024).

54. Chittrakanwong, J. et al. 5-Methyluridine is Ubiquitous in Pseudomonas aeruginosa tRNA and Modulates Antimicrobial Resistance and Virulence. J. Mol. Biol. 437, 169020 (2025).

55. Riquelme-Barrios, S. et al. Epitranscriptomic control of stress adaptations in *Escherichia coli*. Nucleic Acids Res. 54, (2026).

56. Gokhale, N. S. et al. N6-Methyladenosine in Flaviviridae Viral RNA Genomes Regulates Infection. Cell Host Microbe 20, 654–665 (2016).

57. Courtney, D. G. et al. Epitranscriptomic Addition of m5C to HIV-1 Transcripts Regulates Viral Gene Expression. Cell Host Microbe 26, 217–227 (2019).

58. Rodell, R. et al. PUS7 mRNA pseudouridylation is driven by RNA sequence, structure, and cell-type-specific features. Cell Genomics 101332 (2026) doi:10.1016/j.xgen.2026.101332.

59. Begik, O. et al. Quantitative profiling of pseudouridylation dynamics in native RNAs with nanopore sequencing. Nat. Biotechnol. 39, 1278–1291 (2021).

60. Tavakoli, S. et al. Semi-quantitative detection of pseudouridine modifications and type I/II hypermodifications in human mRNAs using direct long-read sequencing. Nat. Commun. 14, 334 (2023).

61. McCormick, C. A. et al. Multicellular, IVT-derived, unmodified human transcriptome for nanopore-direct RNA analysis. Gigabyte 2024, 1–13 (2024).

62. Zhang, M. et al. Quantitative profiling of pseudouridylation landscape in the human transcriptome. Nat. Chem. Biol. 19, 1185–1195 (2023).

63. Dai, Q., et al. Quantitative sequencing using BID-seq uncovers abundant pseudouridines in mammalian mRNA at base resolution. Nat. Biotechnol. 41, 344–354 (2023).

64. Zhang, L.-S. et al. BID-seq for transcriptome-wide quantitative sequencing of mRNA pseudouridine at base resolution. Nat. Protoc. (2023) doi:10.1038/s41596-023-00917-5.

65. Bebenek, K., Abbotts, J., Wilson, S. H. & Kunkel, T. A. Error-prone polymerization by HIV-1 reverse transcriptase. Contribution of template-primer misalignment, miscoding, and termination probability to mutational hot spots. J. Biol. Chem. 268, 10324–10334 (1993).

66. Potapov, V. et al. Discrete measurements of RNA polymerase and reverse transcriptase fidelity reveal evolutionary tuning. RNA 30, 1246–1258 (2024).

67. Shibutani, S., Takeshita, M. & Grollman, A. P. Translesional Synthesis on DNA Templates Containing a Single Abasic Site. J. Biol. Chem. 272, 13916–13922 (1997).

68. Busan, S. & Weeks, K. M. Accurate detection of chemical modifications in RNA by mutational profiling (MaP) with ShapeMapper 2. RNA 24, 143–148 (2018).

69. He, Z., Qiu, W. C Zhou, H. Promoted read-through and mutation against pseudouridine-CMC by an evolved reverse transcriptase. *Commun*. Biol. 8, 40 (2025).

70. Meng, R. et al. Structural basis for the adsorption of a single-stranded RNA bacteriophage. Nat. Commun. 10, 3130 (2019).

71. Chamakura, K. R., Edwards, G. B. & Young, R. Mutational analysis of the MS2 lysis protein L. Microbiology 163, 961–969 (2017).

72. Mezhyrova, J., et al. *In vitro* characterization of the phage lysis protein MS2-L. Microbiome Res. Rep. 2, (2023).

73. Pan, H., Agarwalla, S., Moustakas, D. T., Finer-Moore, J. & Stroud, R. M. Structure of tRNA pseudouridine synthase TruB and its RNA complex: RNA recognition through a combination of rigid docking and induced fit. Proc. Natl. Acad. Sci. 100, 12648–12653 (2003).

74. Carlile, T. M. et al. mRNA structure determines modification by pseudouridine synthase 1. Nat. Chem. Biol. 15, 966–974 (2019).

75. Wrzesinski, J., Nurse, K., Bakin, A., Lane, B. G. & Ofengand, J. A dual-specificity pseudouridine synthase: an Escherichia coli synthase purified and cloned on the basis of its specificity for psi746 in 23S RNA is also specific for psi32 in tRNA(phe). RNA 1, 437–448 (1995).

76. Raychaudhuri, S., Niu, L., Conrad, J., Lane, B. G. & Ofengand, J. Functional Effect of Deletion and Mutation of the Escherichia coli Ribosomal RNA and tRNA Pseudouridine Synthase RluA. J. Biol. Chem. 274, 18880–18886 (1999).

77. Serra, M. J., Lyttle, M. H., Axenson, T. J., Schadt, C. A. & Turner, D. H. RNA hairpin loop stability depends on closing base pair. Nucleic Acids Res. 21, 3845–3849 (1993).

78. Fleming, A. M., Mathewson, N. J., Howpay Manage, S. A. & Burrows, C. J. Nanopore Dwell Time Analysis Permits Sequencing and Conformational Assignment of Pseudouridine in SARS-CoV-2. ACS Cent. Sci. 7, 1707–1717 (2021).

79. Fleming, A. M. & Burrows, C. J. Nanopore sequencing for N1-methylpseudouridine in RNA reveals sequence-dependent discrimination of the modified nucleotide triphosphate during transcription. Nucleic Acids Res. 51, 1914–1926 (2023).

80. Adams, M. Bacteriophages. (Wiley Interscience, New York, 1959).

81. Ellis, E. L. & Delbrück, M. The Growth of Bacteriophage. J. Gen. Physiol. (1937).

82. Abedon, S. Dos and Don’ts of Bacteriophage One-Step Growth. Preprint at 10.20944/preprints202507.2624.v1 (2025).

83. Ni, N. & Burgio, G. Nanopore direct RNA sequencing (DRS) of MS2 bacteriophages in E. coli throughout its life cycles reveals a complex transcriptional activity to control and maintain its growth. Virol. J. 23, 99 (2026).

84. Jones, J. D. et al. Integrating mass spectrometry with Nanopore direct RNA sequencing for de novo modification profiling of bacteriophage MS2.

85. Baek, A. et al. Single-molecule epitranscriptomic analysis of full-length HIV-1 RNAs reveals functional roles of site-specific m6As. Nat. Microbiol. 9, 1340–1355 (2024).

86. Horner, S. M. & Thompson, M. G. Challenges to mapping and defining m^6^ A function in viral RNA. RNA 30, 482–490 (2024).

87. Campos, C. M., et al. Mapping of pseudouridine residues on cellular and viral transcripts using a novel antibody-based technique. (2021).

88. Giambruno, R. et al. Unveiling the role of PUS7-mediated pseudouridylation in host protein interactions specific for the SARS-CoV-2 RNA genome. Mol. Ther. Nucleic Acids 34, 102052 (2023).

89. Xu, H., et al. Absolute Quantitative and Base-Resolution Sequencing Reveals Comprehensive Landscape of Pseudouridine across the Human Transcriptome. http://biorxiv.org/lookup/doi/10.1101/2024.01.08.574649 (2024) doi:10.1101/2024.01.08.574649.

90. Stark, N. et al. Pseudouridine synthases are proviral factors for Sindbis virus in insect and mammalian cells. mBio 16, e01329–25 (2025).

91. Wu, C. C. et al. The two-step purification method ViREn identifies a single NSUN6-mediated 5-methylcytosine modification promoting dengue virus RNA genome turnover. Nucleic Acids Res. 54, (2026).

92. Karikó, K. et al. Incorporation of Pseudouridine Into mRNA Yields Superior Nonimmunogenic Vector With Increased Translational Capacity and Biological Stability. Mol. Ther. 16, 1833–1840 (2008).

93. Eyler, D. E. et al. Pseudouridinylation of mRNA coding sequences alters translation. Proc. Natl. Acad. Sci. 116, 23068–23074 (2019).

94. Olejniczak, M. & Uhlenbeck, O. tRNA residues that have coevolved with their anticodon to ensure uniform and accurate codon recognition. Biochimie 88, 943–950 (2006).

95. Dai, X. et al. In situ structures of the genome and genome-delivery apparatus in a single-stranded RNA virus. Nature 541, 112–116 (2017).

96. Dykeman, E. C., Stockley, P. G. & Twarock, R. Packaging Signals in Two Single-Stranded RNA Viruses Imply a Conserved Assembly Mechanism and Geometry of the Packaged Genome. J. Mol. Biol. 425, 3235–3249 (2013).

97. Rastandeh, A., et al. Measuring the selective packaging of RNA molecules by viral coat proteins in cells. Proc. Natl. Acad. Sci. 122, e2505190122 (2025).

98. Amitsur, M., Levitz, R. & Kaufmann, G. Bacteriophage T4 anticodon nuclease, polynucleotide kinase and RNA ligase reprocess the host lysine tRNA. EMBO J. 6, 2499–2503 (1987).

99. Hazan, R. & Engelberg-Kulka, H. Escherichia coli mazEF-mediated cell death as a defense mechanism that inhibits the spread of phage P1. Mol. Genet. Genomics 272, 227–234 (2004).

100. Klaiman, D., Steinfels-Kohn, E., Krutkina, E., Davidov, E. & Kaufmann, G. The wobble nucleotide-excising anticodon nuclease RloC is governed by the zinc-hook and DNA-dependent ATPase of its Rad50-like region. Nucleic Acids Res. 40, 8568–8578 (2012).

101. Belshaw, R., Pybus, O. G. & Rambaut, A. The evolution of genome compression and genomic novelty in RNA viruses. Genome Res. 17, 1496–1504 (2007).

102. Zhang, T. et al. Bacterial immune activation via supramolecular assembly with phage triggers. Nature 651, 1051–1059 (2026).

103. Winther, K. S., Brodersen, D. E., Brown, A. K. & Gerdes, K. VapC20 of Mycobacterium tuberculosis cleaves the Sarcin–Ricin loop of 23S rRNA. Nat. Commun. 4, 2796 (2013).

104. Baba, T. et al. Construction of Escherichia coli K-12 in-frame, single-gene knockout mutants: the Keio collection. Mol. Syst. Biol. 2, 2006.0008 (2006).

105. Elston, K. M. et al. The Pathfinder plasmid toolkit for genetically engineering newly isolated bacteria enables the study of *Drosophila*-colonizing *Orbaceae*. ISME Commun. 3, 49 (2023).

106. Small, E. et al. 1 A reference genome without a virus: cDNA reconstruction reveals the 2 provenance and function of the MS2 phage sequence. (2026).

107. Vo, J. M. et al. Synthesis of modified nucleotide polymers by the poly(U) polymerase Cid1: application to direct RNA sequencing on nanopores.

108. Arribere, J. A. et al. Translation readthrough mitigation. Nature 534, 719–723 (2016).

109. Kim, H. et al. Bias-minimized quantification of microRNA reveals widespread alternative processing and 3′ end modification. Nucleic Acids Res. 47, 2630–2640 (2019).

110. Galls, D., Mueller, A. U., Greenwald, E. & Fire, A. A general RNA-templated RNA extension activity of E. coli RNA polymerase. RNA rna.080238.124 (2025) doi:10.1261/rna.080238.124.

111. Letunic, I. & Bork, P. Interactive Tree of Life (iTOL) v6: recent updates to the phylogenetic tree display and annotation tool. Nucleic Acids Res. 52, W78–W82 (2024).

